# Dissection of multiple sclerosis genetics identifies B and CD4+ T cells as driver cell subsets

**DOI:** 10.1101/2021.05.24.445445

**Authors:** Michael H. Guo, Prashanth Sama, Brenna A. LaBarre, Hrishikesh Lokhande, John Balibalos, Ci Chu, Xiaomi Du, Pouya Kheradpour, Charles C Kim, Taylor Oniskey, Thomas Snyder, Damien Z Soghoian, Howard L. Weiner, Tanuja Chitnis, Nikolaos A. Patsopoulos

## Abstract

Multiple sclerosis (MS) is an autoimmune condition of the central nervous system with a well-characterized genetic background. Prior analyses of MS genetics have identified broad enrichments across peripheral immune cells, yet the driver immune subsets are unclear. We utilized chromatin accessibility data across hematopoietic cells to identify cell type-specific enrichments of MS genetic signals. We found that CD4 T and B cells were independently enriched for MS genetics and further refined the driver subsets to T_h_17 and memory B cells, respectively. We replicated our findings in data from untreated and treated MS patients and found that immunomodulatory treatments suppress chromatin accessibility at driver cell types. Integration of statistical fine-mapping and chromatin interactions nominated numerous putative causal genes, illustrating complex interplay between shared and cell-specific genes. Our study highlights how careful integration of genetics and epigenetics can provide fine-scale insights into causal cell types and nominate new genes and pathways for disease.

## Introduction

Multiple sclerosis (MS) is an immune-mediated neurodegenerative disease characterized by demyelinating focal lesions in the central nervous system (CNS)^1^. Despite the CNS being the target of autoimmunity, there is extensive evidence from basic science models and human studies that dysregulation of the peripheral immune compartment is key for disease manifestation and progression ^2^. MS has long been regarded as a T cell-mediated disease, with several T cell subpopulations implicated ^3, 4^. More recently, other peripheral immune cell populations, most notably B cells, have also been shown to drive disease pathogenesis^3, 5^. Moreover, immune modulating therapies targeting B cells have been demonstrated to be remarkably effective in treating patients with MS^6, 7^.

MS has a strong genetic component and is characterized by a polygenic architecture. To date, over 200 independent genetic variants have been associated with MS risk, the vast majority of which are common variants with small effect sizes on disease risk^8, 9^. Prior studies have shown enrichment of GWAS target genes in the peripheral immune system, but it is unclear exactly which cell types within the peripheral immune system drive these observed enrichments of genetic signals.

Human genetics has emerged as a powerful tool for probing the underlying biology of a disease^10^. The identification of genes and pathways prioritized by GWAS associations is not constrained by our prior knowledge of disease mechanisms and can therefore identify novel biological mechanisms. However, a key challenge for translating GWAS findings into biological insights is that most associations are noncoding in nature and likely act by modulating regulatory elements to mediate gene expression^11, 12^. Identifying the causal gene at these GWAS signals can be challenging since it is usually unclear which gene a given regulatory element regulates ^10, 13, 14^. A key step in translating genetic associations into biological mechanisms is identifying the cell types in which GWAS variants act and the genes they modulate. To help understand the function of disease-associated variants, genetic associations can be intersected with orthogonal epigenetic and gene expression data^12^. As the epigenetic and gene expression landscape differ from cell type to cell type, examining the enrichments of GWAS data on these orthogonal datasets can identify specific cell types that may be implicated in disease pathogenesis^15, 16^.

We and others have previously reported a strong enrichment of MS GWAS variants in regulatory regions of multiple cell types of the peripheral immune system^8, 9, 15, 17^. However, it has yet to be determined if these enrichments are driven by shared regulatory mechanisms common to many immune cell types, or whether different mechanisms are present in distinct immune cell populations. To address this gap, we performed detailed analyses of the enrichment of MS GWAS variants in the peripheral immune system to identify cell populations that independently mediate the effects of MS GWAS variants on disease risk.

## Results

### MS GWAS associations are enriched in progenitor and terminal peripheral immune cells

To identify the causal cell types that uniquely and independently contribute to MS pathogenesis via mediation of genetic effects, we leveraged bulk ATAC-seq data from 16 flow-sorted hematopoietic progenitor and terminal cell populations isolated from human peripheral blood or bone marrow^18–20^. These cells represent progenitor and terminal populations from across the hematopoietic tree, enabling investigation of MS GWAS enrichments broadly and across stem, progenitor, and mature cell populations (**Figure 1A**). ATAC-seq data were processed and open chromatin regions (OCRs, i.e. ATAC-seq peaks), were identified as previously described^18, 21^. We applied stratified LD SCore regression (LDSC)^22, 23^ to estimate the enrichment of MS GWAS in OCRs from each of the 16 hematopoietic cell types. LDSC has the distinct advantage in that it leverages the genome-wide polygenic signal in the GWAS summary statistics rather than selecting variants based on p-value thresholds or fine-mapping posterior probabilities.

**Figure 1:**
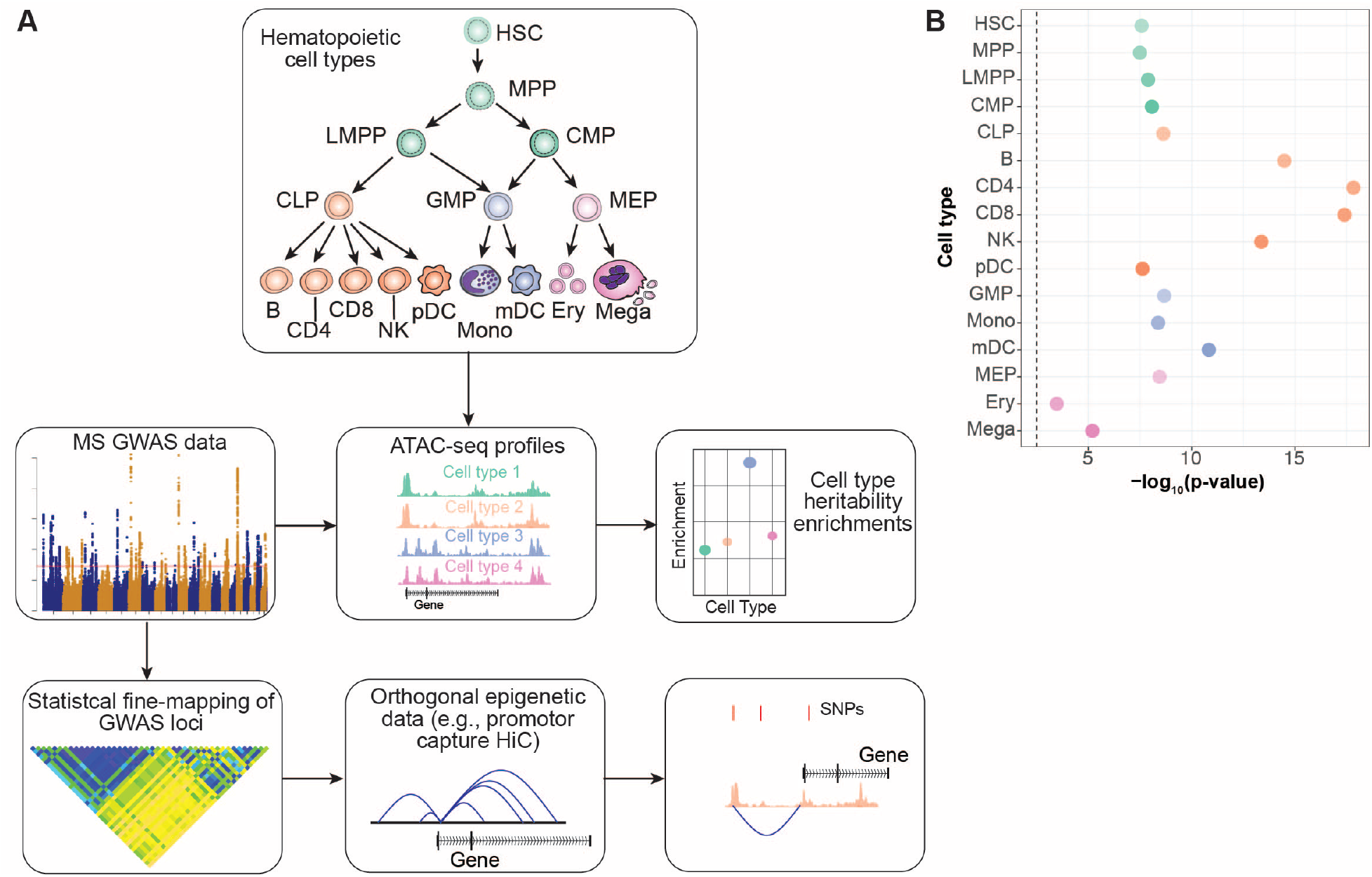
**A:** Experimental Design. Top box shows the hematopoietic cell types analyzed. MS discovery GWAS results were integrated with ATAC-seq profiles generated from the hematopoietic cell types. LDSC was performed to evaluate enrichment of MS GWAS in the OCRs of each hematopoietic cell type. Statistical fine-mapping was also performed on the MS GWAS results, which were then integrated with orthogonal epigenetic data such as promotor capture HiC interactions. This integration of fine-mapping and epigenetic data allowed for identification of putative causal mechanisms at individual loci. **B:** Enrichment of MS GWAS heritability in hematopoietic cell OCRs. Enrichment p-values are shown as –log_10_(p-value).

Applying LDSC, we observed strong statistically significant enrichments across all hematopoietic cell populations, even after correcting for multiple hypothesis testing (Bonferroni-corrected p-value threshold of 3.13x10^-3^) (**Figure 1B; Supplemental Table 1**). The strongest enrichments for MS GWAS were observed in OCRs from CD4 T cells (enrichment p-value = 1.47x10^-18^), CD8 T cells (p-value = 4.00 x10^-18^) and B cells (p-value = 3.27x10^-15^); reflecting their known and emerging roles in MS pathogenesis and as targets of treatment^1, 3, 24^. We also detected strong enrichment in OCRs from natural killer (NK) cells (p-value =4.23x10^-14^) which have a less well-established role in MS^24^. Interestingly, we observed enrichments across all progenitor cells, suggesting that many MS genetic associations are located in regulatory regions involved in core cellular processes in immune cell populations.

### CD4 T and B cell regulatory regions independently mediate MS genetics

Many of the studied cell populations share common cellular regulatory signatures, which is reflected in the substantial correlation of the OCR profiles across cell populations (**Figure S1**). Hence, we examined whether the strong enrichment observed across cell populations is a result of truly independent cell type-specific enrichments or whether it is due to shared regulatory landscape across immune cell types. To address this question, we applied a joint model in LDSC to measure the contribution of OCRs from a given cell type, stratified on all other cell types in the model along with a set of baseline annotations. We report the p-value of the coefficient *τ_c_*, which reflects the SNP heritability of a given annotation stratified on all other annotations in the model. In this joint model where OCRs from all 16 cell types were included, we observed that B cells and CD4 T cells contributed significantly to SNP heritability (coefficient p-value = 3.99x10_-5_ and 3.49x10^-4^, respectively), suggesting independent contributions of B and CD4 T cell OCRs to MS GWAS heritability (**Figure 2A, Supplemental Table 2**).

**Figure 2:**
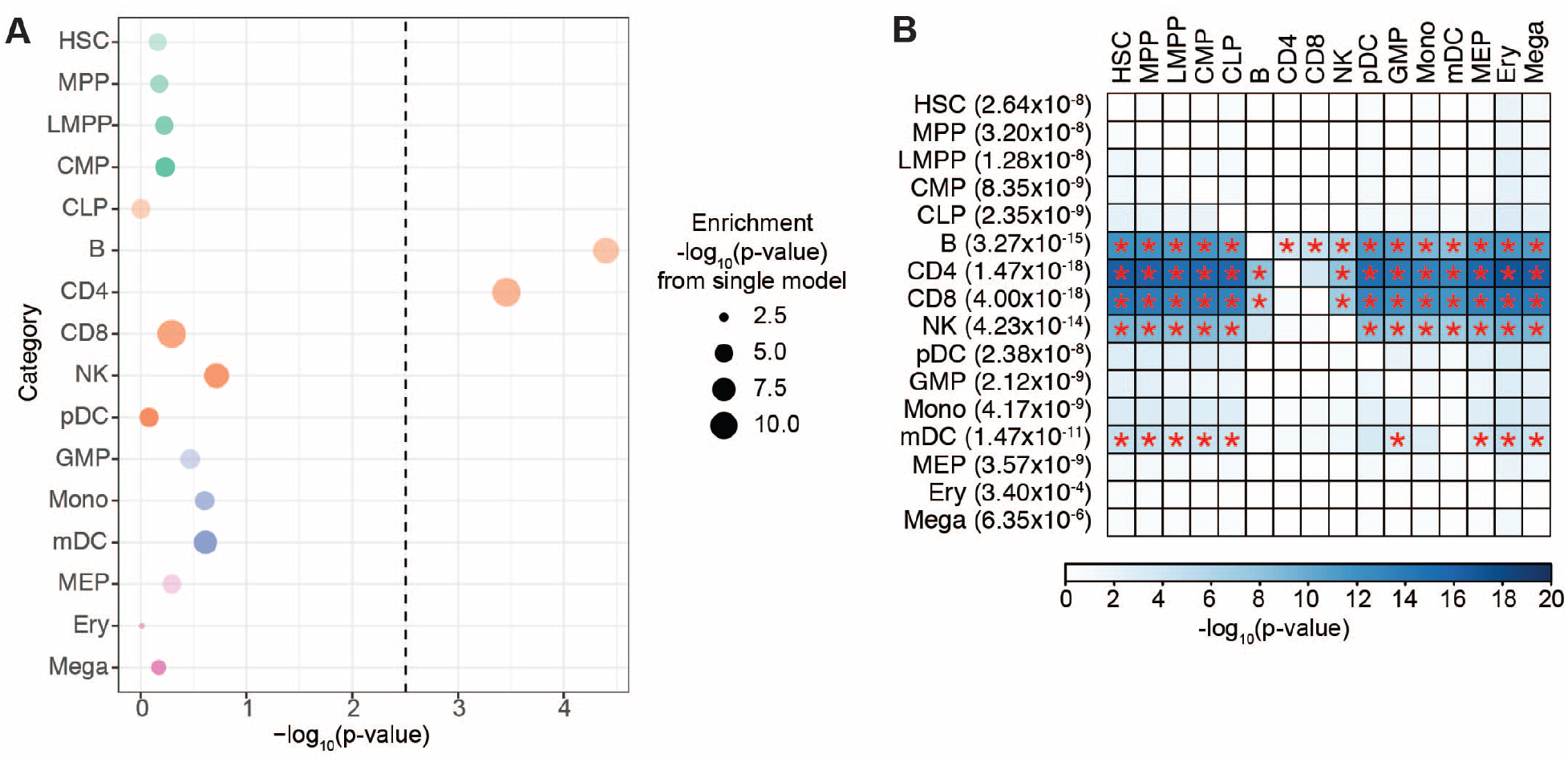
**A:** LDSC enrichment results for MS GWAS enrichment in OCRs from across hematopoietic cell types in a joint model. Heights of the circles reflect LDSC coefficient (*τ_c_*) p-values, which measures whether the annotation (i.e., OCRs for a given cell type) contributes significantly to SNP heritability in an overall model that includes OCRs for all hematopoietic cell types and baseline annotations. Sizes of the circles are proportional to the enrichment p-values for that given cell type, with larger circles reflecting more significant p-values. **B:** LDSC enrichment p-values for pairwise stratified LDSC of MS GWAS results in OCRs from hematopoietic cell types. Y-axis are the index cell types with LDSC enrichment p-values prior to stratifying in parentheses. X-axis shows the comparator cell type being conditioned upon. Boxes are shaded by the LDSC coefficient p-values for the index cell type after conditioning on the comparator cell type in the pairwise model (with darker colors representing stronger enrichments). Red stars indicate pair-wise comparisons that are statistical significant a Bonferroni corrected p-value threshold of 2.2x10^-4^.

To further delineate the cell types with OCRs that specifically mediate the effect of MS GWAS results, we performed a series of pairwise LDSC analyses. In brief, OCRs of a given hematopoietic cell type were stratified against the OCRs of each of the other 15 cell types, as well as the LDSC baseline annotations (**Figure 2B; Supplemental Table 3**). As above with the joint model, we report the p-value of the coefficient *τ_c_*. We observed that B cells remained significant even after stratifying on OCRs of any of the other 15 cell populations (coefficient p-value ranging from 1.47x10^-12^ when stratifying against HSCs, to 8.33x10^-5^ when stratifying against CD4 T cells). This was also the case for CD4 T cells, which remained significant after stratifying on OCRs from any of the other 15 cell populations (coefficient p-values ranging from 3.82x10^-17^ when stratifying against HSCs, to 2.39x10^-4^ when stratifying against CD8 T cells). In contrast, CD8 T cell OCRs were no longer significant after stratifying against OCRs from CD4^+^ T cells (coefficient p-value = 0.21), though they were significant when stratifying on any of the other cell populations. NK cells were also no longer significant after stratifying against either CD4 T cells (coefficient p-value = 0.165) or against CD8 T cells (coefficient p-value: 0.356). These results indicate that the enrichment of both CD8 T cells and NK cells can be largely explained by shared regulatory landscapes that are also present in CD4 T cells. Prior studies have also suggested a role for monocytes in MS^24, 25^. OCRs from monocytes had an enrichment p-value of 4.17x10^-9^, but we observed that stratifying on OCRs from CD4 T cells or B cells ameliorated this monocyte heritability enrichment (coefficient p-values 0.166 and 0.266, respectively).

We also performed a separate analysis examining whether OCRs specific to a given cell type mediate MS heritability enrichments. For each of the mature hematopoietic cell populations, we performed LDSC on only the cell type-specific OCRs, i.e. ATAC-seq peaks present in only that cell type. B cells exhibited a statistically significant enrichment of cell-specific OCRs for MS GWAS (enrichment p-value = 3.27x10^-4^). CD4 T cell-specific peaks were nominally significant at a p-value of 0.013 but did not survive correction for multiple hypothesis testing (Bonferroni corrected p-value threshold of 5.6x10^-3^). Cell type-specific peaks for all other terminal hematopoietic cell types had enrichment p-values > 0.05 (**Figure S2; Supplementary Table 4**).

### MS genetic associations are mediated in terminal immune cell populations

In the lymphoid lineage, we observed stronger enrichments in terminal cell populations than we did for the progenitor populations (**Figure 1B**). For example, the strongest enrichment in progenitor cells was observed for common lymphoid progenitor cells (CLP, enrichment p-value: 2.32x10^-4^), and it was orders of magnitude less statistically significant compared to the enrichment observed for CD4 T or B cells (**Figure 1B**). For each of the terminal populations, the significance remained even after stratifying against OCRs from CLP; however, the converse was not true (**Figure 2B**). The enrichment in CLP was completely ameliorated by stratifying against B cell or CD4 T cell OCRs (coefficient p-value 0.98 and 0.966, respectively, **Figure 2B**). Together, these results suggest that terminal cells of the lymphoid compartment retain cellular regulatory features from their progenitor populations that are important for MS pathogenesis, but have also developed specific regulatory features of additional importance to MS susceptibility.

### Comparison of immune cell enrichment with neuropsychiatric and autoimmune disorders

We next sought to understand how the immune cell enrichments in MS might be similar or different from those of other autoimmune or neuropsychiatric disorders. To test this, we calculated heritability enrichment within these 16 hematopoietic OCRs for GWAS of various other neuropsychiatric disorders and autoimmune disorders: Alzheimer disease (AD)^26^, schizophrenia (SCZ)^27^, bipolar disorder (BPD)^28^, type 1 diabetes (T1D)^29^, Crohn’s disease (CD)^30^, ulcerative colitis (UC)^30^, systemic lupus erythematosus (SLE)^31^, rheumatoid arthritis (RA)^32^, and primary biliary cirrhosis (PBC)^33^ (**Figure 3A; Supplemental Table 5**). We identified previously recognized cell-type enrichments across these other diseases, such as enrichments in OCRs from B cells (enrichment p-value: 1.93x10^-7^), CD4 T cells (p-value: 7.34x10^-8^) and CD8 T cells (p-value: 2.09x10^-6^) for RA^15, 32^. However, the heritability enrichments for OCRs from these hematopoietic cell populations tended to be much stronger for MS, despite similar sample sizes, e.g. 41,505 disease cases for MS and 38,242 disease cases for RA. To parse out cell type-specific enrichments in these other disorders, we also tested heritability enrichments under the joint model in LDSC by including OCRs from all 16 cell types (**Figure 3B, Supplemental Table 6**). In a similar fashion to the MS GWAS joint analyses above, we measured the contribution to heritability for a given set of OCRs of interest, controlling for the effects of a set of baseline annotations and OCRs from all other hematopoietic cell types. These analyses were performed separately for each disease. Across the nine other comparator diseases, the only other statistically significant stratified enrichments were in OCRs from B cells in SLE GWAS (coefficient p-value: 8.53x10^-5^). We also identified other enrichments in this joint model that were nominally significant, including signals for several diseases in either B or CD4 T cells. This is in contrast to our MS findings, which show strong enrichments in both CD4 T cells and B cells, highlighting a dual role of these cell types in MS and distinguishing it from other diseases where the cell types are more restricted to specific lineages.

**Figure 3:**
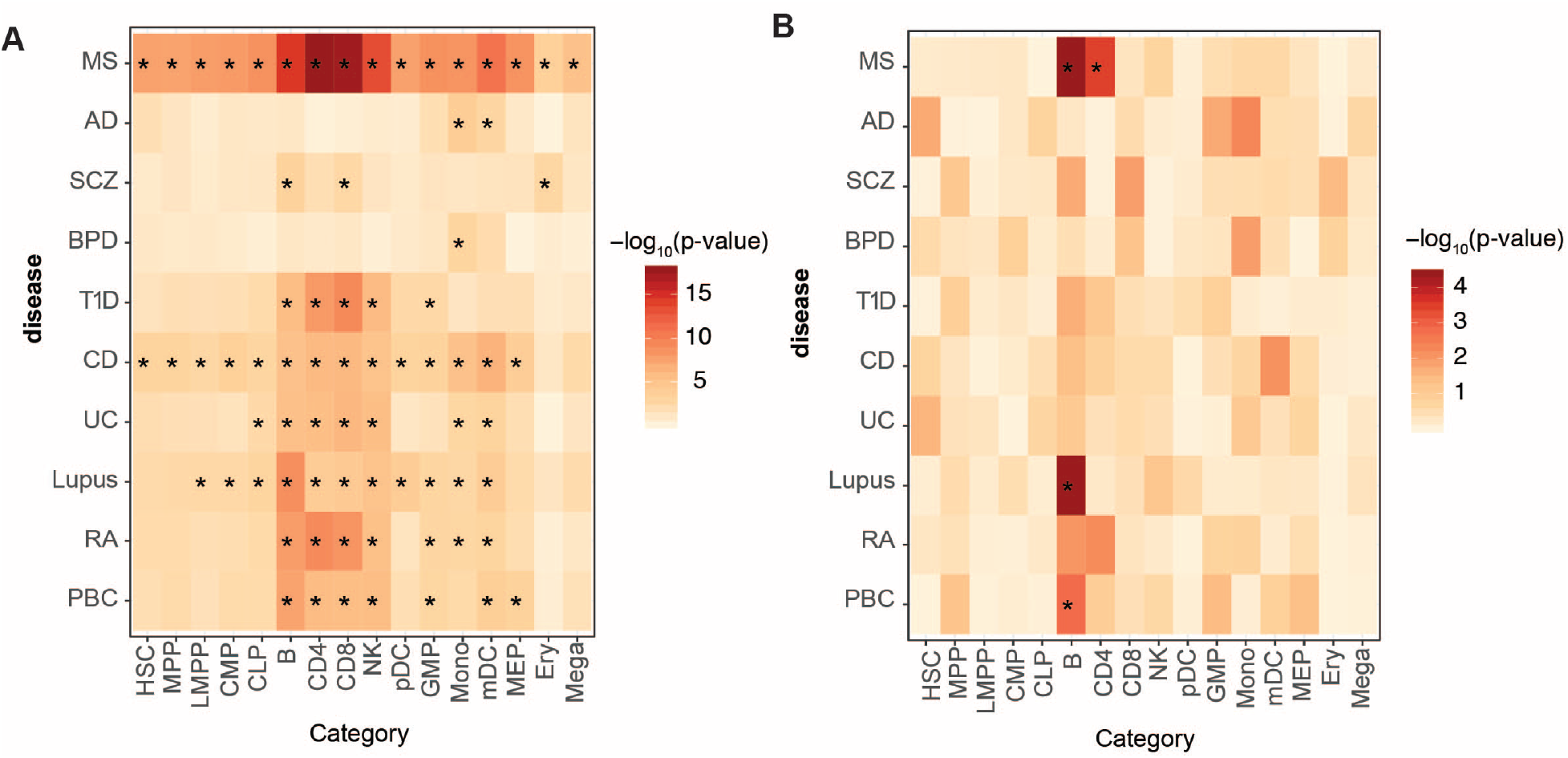
**A:** Enrichments of GWAS results from 10 neuropsychiatric or autoimmune conditions in OCRs across various hematopoietic cell types. **B:** LDSC coefficient p-values in the joint model across hematopoietic cell ATAC-seq in 10 neuropsychiatric or autoimmune conditions. For **A** and **B**, boxes are shaded by –log_10_(p-value), with darker shading reflecting more statistical significance, and statistically significant p-values (p-values<3.13x10^-3^) are starred.

### The CD4 T cell MS GWAS enrichment is driven by the Th17 subset

To further focus in on the MS GWAS enrichment observed in OCRs from CD4 T cells, we utilized ATAC-seq data from various subsets of human CD4 T cells (**Figure 4A**)^21^. We examined data from effector CD4 T cells (naïve effector CD4 T cells, T_h_1, T_h_2, T_h_17, and follicular T_h_) as well as, regulatory CD4 T cells (naïve T_regs_ and memory T_regs_). We observed strong enrichments for OCRs from both effector and regulatory CD4 T cell populations (**Figure 4B; Supplementary Table 7**). Next, to identify the independent contribution of a given cell type, we applied the joint model in LDSC by including OCRs from all CD4 T cell populations together. This joint LDSC analysis revealed that OCRs from T_h_17 cells independently contributed to heritability (coefficient p-value = 4.69x10^-4^; **Figure 4C; Supplemental Table 8**). Performing pairwise stratified LDSC confirmed the independent contribution of OCRs in T_h_17 cells to MS GWAS heritability **(Figure 4D; Supplemental Table 9**). OCRs from T_h_17 remained enriched in MS GWAS even after stratifying against any of the other T_eff_ cell populations or T_reg_ cell populations. Conversely, the enrichments for all other T_eff_ cell populations or T_reg_ cell populations were ameliorated when stratifying against T_h_17 cells.

**Figure 4:**
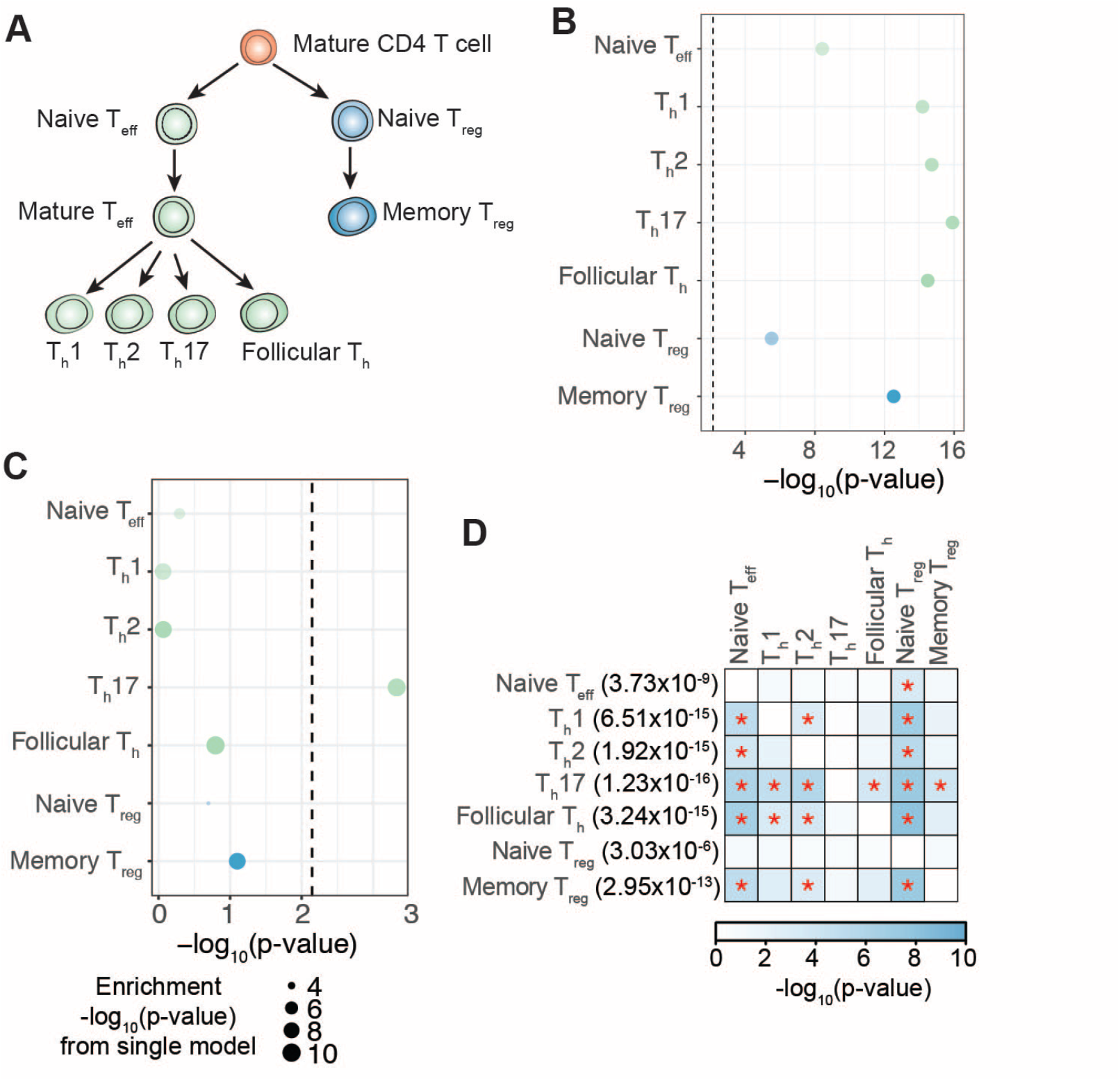
**A:** Schematic of lineage relationships among CD4^+^ T cell subsets for which ATAC-seq data was analyzed. **B:** LDSC heritability enrichment p-values for CD4^+^ T cell subsets in MS GWAS. See Figure 1B for additional description. **C:** LDSC coefficient p-values for CD4^+^ T cells in MS GWAS. See Figure 2A for additional description. **D:** LDSC coefficient p-values for pairwise stratified analyses of MS GWAS results in ATAC-seq data from CD4^+^ T cell subsets. See Figure 2B legend for additional description.

Based on the LDSC analyses in the joint model, we note that enrichments tended to be stronger in OCRs from the memory effector CD4 T cell populations than from naïve effector cells. For each of these memory effector CD4 T cell populations, OCRs retained statistical significance even after stratifying against OCRs from naïve T_eff_ cells: T_h_1 (coefficient p-value after stratifying against naïve T_eff_ cells: 8.22x10^-6^), T_h_2 (p-value = 1.45x10^-4^), T_h_17 (p-value = 7.27x10^-7^) or follicular T_h_ cells (p-value = 1.77x10^-7^) (**Figure 4D**). The converse was not true; the statistical significance of the naïve T_eff_ cell OCRs was completely lost when stratifying against OCRs from any of the memory T_eff_ cell populations. These results suggest that among CD4 T cells, OCRs from T_h_17 cells drive the signal for enrichment in MS GWAS.

### Memory subpopulations explain the enrichment of MS GWAS in B cells

We next examined enrichments of MS GWAS data in ATAC-seq from the B cell lineage including naïve B cells, memory B cells, and plasmablasts (**Figure 5A**) ^21^. We found that OCRs from all B cell lineage cell types were significantly enriched for MS heritability (**Figure 5B; Supplemental Table 10**). LDSC under a joint model including all B cell lineage cell types revealed an independent contribution from memory B cells (coefficient p-value = 1.10 x 10^-3^), but not naïve B cells or plasmablasts **Figure 5C; Supplemental Table 11**). Pairwise stratified LDSC confirmed the independent enrichment of OCRs from memory B cells. Memory B cell OCRs remained statistically significant even after stratifying on naïve B cells (coefficient p-value = 1.26x10^-5^) or plasmablasts (coefficient p-value: 3.29x10^-4^; **Figure 5D; Supplemental Table 12**). In contrast, OCRs from naïve B cells and plasmablasts were no longer statistically significant when stratifying on memory B cells OCRs (coefficient p-value: 0.71 and 0.27, respectively). These results suggest that in the B cell lineage, the MS GWAS enrichment signal is driven by OCRs in memory B cells.

**Figure 5:**
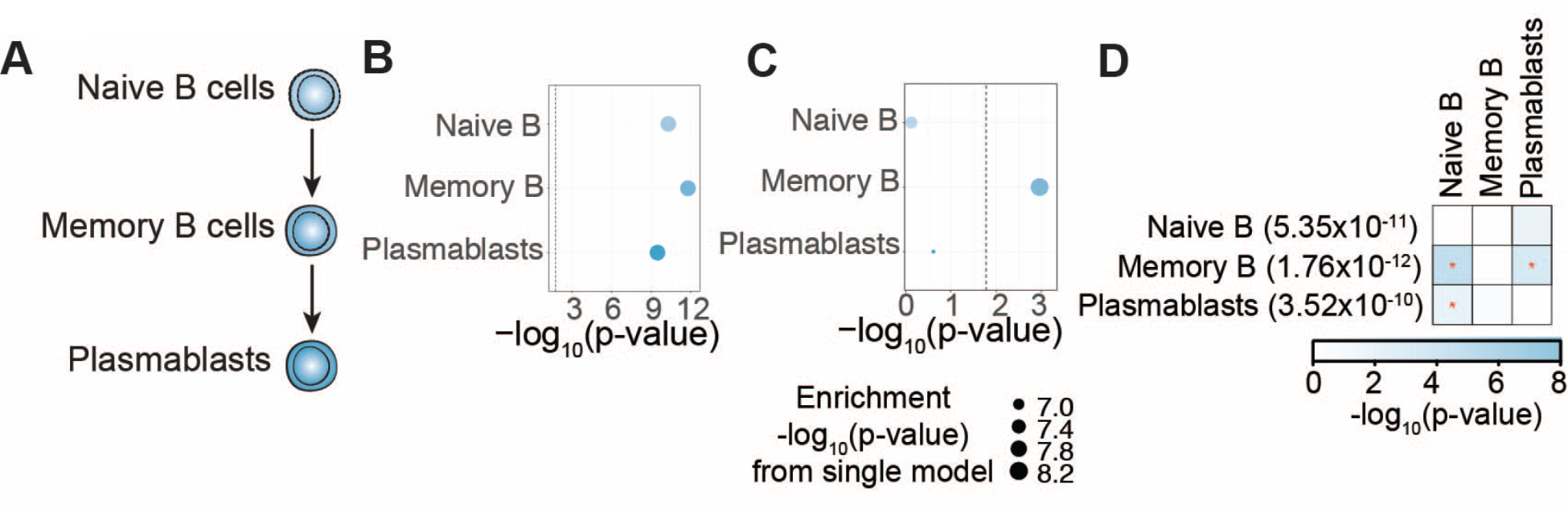
**A:** Schematic of lineage relationships among B cell lineage cell types for which ATAC-seq data was analyzed. **B:** LDSC heritability enrichment p-values for B cell lineage cell types in MS GWAS. See Figure 1B for additional description. **C:** Stratified LDSC coefficient p-values for B cell lineage cell types in MS GWAS. See Figure 2A for additional description. **D:** LDSC coefficient p-values for pairwise stratified analyses of MS GWAS results in ATAC-seq data from B cell lineage cell types. See Figure 2B legend for additional description.

### Immune cells from MS patients reinforce independent CD4 T and B cell enrichments

Next, we tested whether the MS GWAS enrichment in CD4+ T and B cells were also present in OCRs from the respective immune cell types derived from individuals with MS. We utilized ATAC-seq data performed in flow-sorted bulk CD4 T and B cell subsets (n=6) derived from six patients with MS who were not treated with immunomodulatory therapy within at least 4 months of sample collection (see **Supplementary Table 13** for clinical details). LDSC showed statistically significant enrichments of transitional B cells (traB; enrichment p-value=2.08x10^-3^), class switched classical memory B cells (cMBc; p-value = 2.58x10^-4^), effector memory CD4 T cells (T4_em_; p-value = 1.87x10^-4^), and CD45RA+ effector memory CD4 T cells (T4ra; p-value = 3.02x10^-4^). Central memory CD4 T cells had an enrichment p-value of 1.53x10^-3^, which was not significant after correcting for multiple hypothesis testing (**Figure 6A; Supplementary Table 14**).

**Figure 6:**
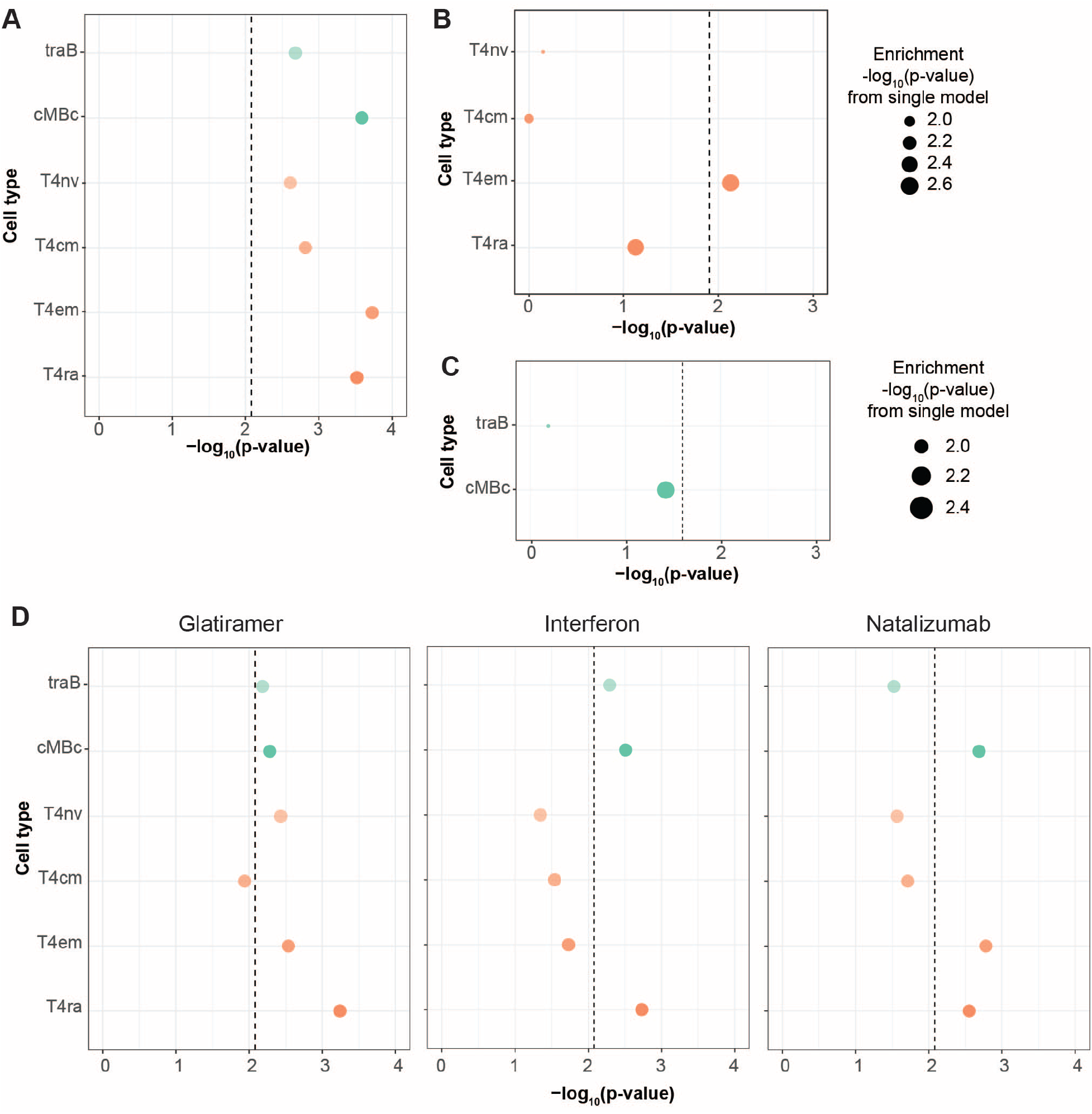
**A:** Enrichment of MS GWAS heritability in OCRs from untreated patients with MS. Enrichment p-values are shown as –log_10_(p-value). **B, C:** LDSC results for MS GWAS enrichment in a joint model for T4cm (**B**) and cMBc (**C**) OCRs from untreated patients with MS. Heights of the circles reflect stratified LDSC coefficient p-values. Sizes of the circles are proportional to the enrichment p-values for that given cell type, with larger circles reflecting more significant p-values. **D:** Enrichment of MS GWAS heritability in OCRs from MS patients undergoing immunomodulatory treatment. Enrichment p-values are shown as –log_10_(p-value). Treatments include glatiramer acetate, interferon, or natalizumab.

To further identify independent cell type enrichments, we performed joint models in LDSC as described above. We first tested a model that included CD4 T cell subsets (T4nv, T4cm, T4em, and T4ra). In this joint model, only T4em was statistically significant (coefficient p-value 7.45 x 10^-3^), reflecting the independent contribution of effector CD4 T cells in MS that we observed above using cells from healthy individuals (**Figure 6B; Supplementary Table 15**). We also ran a model that included B cell subsets (traB and cMBc). In this joint comparison, neither cell type was statistically significant when correcting for multiple hypothesis testing. However, cMBc had a coefficient p-value that was nominally significant (p-value=0.037), reiterating the independent contributions of mature B cell types (**Figure 6C; Supplementary Table 16**).

### Immunomodulatory treatments suppress mediation of MS genetics in cell-specific fashion

Next, we tested whether immunomodulatory treatments alter the cell-specific mediation of MS genetic associations by utilizing data for the same immune subsets sorted from patients with MS (n=3) under treatment with either natalizumab, interferon, or glatiramer acetate. Following treatment with any of the agents still resulted in statistically significant enrichments of cMBc and T4em (**Figure 6D; Supplementary Table 17**), though the magnitude of the enrichments were attenuated relative to the signals observed from cells from untreated patients (compare **Figure 6A** **and 6D).** To better understand this attenuation of enrichments, we ran joint models in LDSC for T4em and cBMc cells in which we included OCRs from untreated patients and treated patients. In a joint model with T4em OCRs from treated and untreated MS patients, only OCRs from untreated patients were statistically significant (**Figure S3A; Supplementary Tables 18**). Similarly, in a joint model with cMBc OCRs from treated and untreated MS patients, only OCRs from untreated patients were statistically significant (**Figure S3B; Supplementary Table 19**). Together, these results suggest that immune-modulating therapies may attenuate the chromatin accessibility signals at MS GWAS.

### MS GWAS signals in B and CD4 T cells driven by active enhancer and promoter regions

We next sought to gain further insight into the functional consequences of the B and CD4 T cell OCRs underlying MS GWAS signals. We examined enrichments for MS GWAS in chromatin immunoprecipitation sequencing (ChIP-seq) peaks from various histone modifications (H3K27ac, H3K27me3, H3K36me3, H3K4me1, H3K4me3, and H3K9me3) ^34, 35^. In T_h_17 cells (**Figure S4A; Supplementary Table 20**), we detected statistically significant enrichments for H3K27ac (enrichment p-value = 6.54x10^-9^), H3K4me1 (p-value = 3.96x10^-19^) and H3K4me3 (p-value = 2.29x10^-8^). Similarly, in B cells (**Figure S4B; Supplementary Table 21**), we also detected significant enrichments for H3K27ac (enrichment p-value = 1.18x10^-11^), H3K4me1 (p-value = 4.96x10^-16^) and H3K4me3 (p-value = 3.93x10^-8^). These results suggest that MS genetic association are primarily enriched at active noncoding elements: primed enhancers (H3K4me1), active enhancers (H3K27ac and H3K4me1) and active promoters (H3K4me3).

To further delineate the chromatin states with the strongest MS genetic associations, we examined enrichments of MS GWAS results in the predicted chromatin states for B cells and T_h_17 T cells as available in the RoadMap Epigenomics Project ^36^. For Th17 CD4 T cells, the “EnhA2” chromatin state (Active Enhancer 2) were statistically enriched (enrichment p-value=1.19 x 10^-3^) (**Figure S5A; Supplementary Table 22)**. For B cells, “Tx3” (Transcribed 3’ preferential; enrichment p-value=1.80x10^-3^) and “PromD1” (Promoter Downstream TSS 1; p-value=4.40x10^-4^) were statistically enriched, again reflecting the strongest enrichments at active regulatory elements (**Figure S5B; Supplementary Table 23)**. These results demonstrate that MS GWAS variants act through activating regulatory elements, consistent with the autoimmune nature of MS.

### Fine-mapping of MS GWAS loci in cell-specific OCRs

We next sought to understand underlying mechanisms by nominating putative causal genes and variants in a cell-specific fashion. First, we applied statistical fine-mapping to nominate likely causal SNPs. Most statistical fine-mapping approaches require association information across all SNPs in a given locus. In contrast, for MS GWAS, genome-wide results are based on targeted replication analyses, which by design included only a select subset of SNPs within each locus. To overcome this challenge, we applied PICS to perform statistical fine-mapping, which as compared to other statistical fine-mapping approaches, does not require GWAS summary statistics for all SNPs in a region^15^. We defined for each locus a 95% credible set such that the sum of the posterior probabilities for variants in that credible set is greater than or equal to 95%. For fine-mapping, we used the joint analysis MS GWAS results, which include only replicated genome-wide effects^8^. We included all 200 non-MHC loci where the MS GWAS joint association p-value was less than 5 x10^-8^. Next, we prioritized SNPs if they had a PICS posterior probability (PP) > 1% and were included in the 95% PICS credible set. This is a liberal threshold for defining prioritized SNPs, aimed at increasing sensitivity for detecting possible causal variants. Across the 200 loci, there were 3436 credible set variants (1-58 variants per locus) (**Figure S6A**). At 19 loci, there was only one variant in the credible set, and at 37 loci, there were four or fewer prioritized variants (**Figure S6B)**.

Next, we intersected the 3436 credible set variants with OCRs from the 16 hematopoietic cell populations, which we chose to use as they cover a broad range of hematopoietic cell types^18^. Across the 200 loci, 870 of the prioritized variants overlapped an OCR in at least one cell type. Remarkably, 163 out of the 200 loci had at least one prioritized variant overlapping an OCR in any cell type. B and CD4 T cells were the cell types with the greatest number of loci with a credible set SNP overlapping an OCR,126 and 125 loci respectively (**Figure S7**).

### Integration of genetic and epigenetic data identifies putative causal genes

The vast majority of GWAS-associated variants are noncoding and regulate genes that may be far from the variant in terms of linear distance on the genome. To address this challenge and nominate putative causal genes that are regulated by the MS-associated OCRs, we leveraged promoter capture Hi-C data (PCHiC), which identifies chromatin looping interactions between regulatory elements and target gene promoters. We utilized PCHiC data performed on 17 hematopoietic cell populations, which partially overlap with the cell types for which we have ATAC-seq data^37^. These cell populations include naïve B cells, total B cells, activated CD4 T cells, non-activated CD4 T cells, and total T cells. We considered only GWAS loci where a credible set MS GWAS SNP overlapped both an OCR and a PCHiC interaction.

Through these analyses, we nominated 261 genes within 86 MS loci in B cells and 364 genes within 115 MS loci in CD4 T cells (**Supplemental Table 24).** We note that these genes are putative causal genes, representing a list of genes that could be linked with the MS loci via a regulatory mechanism in the respective cell types. The majority of these genes were shared between B and CD4 T cells (n=178; 68.2% and 48.9% respectively; **Figure 7A**). We have previously suggested a list of putative causal genes (n=551) based on an ensemble of methods that did not include ATAC-seq or chromatin interactions^8^. Of these 551 previously nominated genes, 67 (12.2%) overlapped with our B cell prioritized genes and 111 (20.1%) with our CD4 T cell prioritized genes, highlighting that our current mechanism-specific gene prioritization is capturing a large number of potentially causal genes that have not been previously implicated in MS genetic studies.

**Figure 7:**
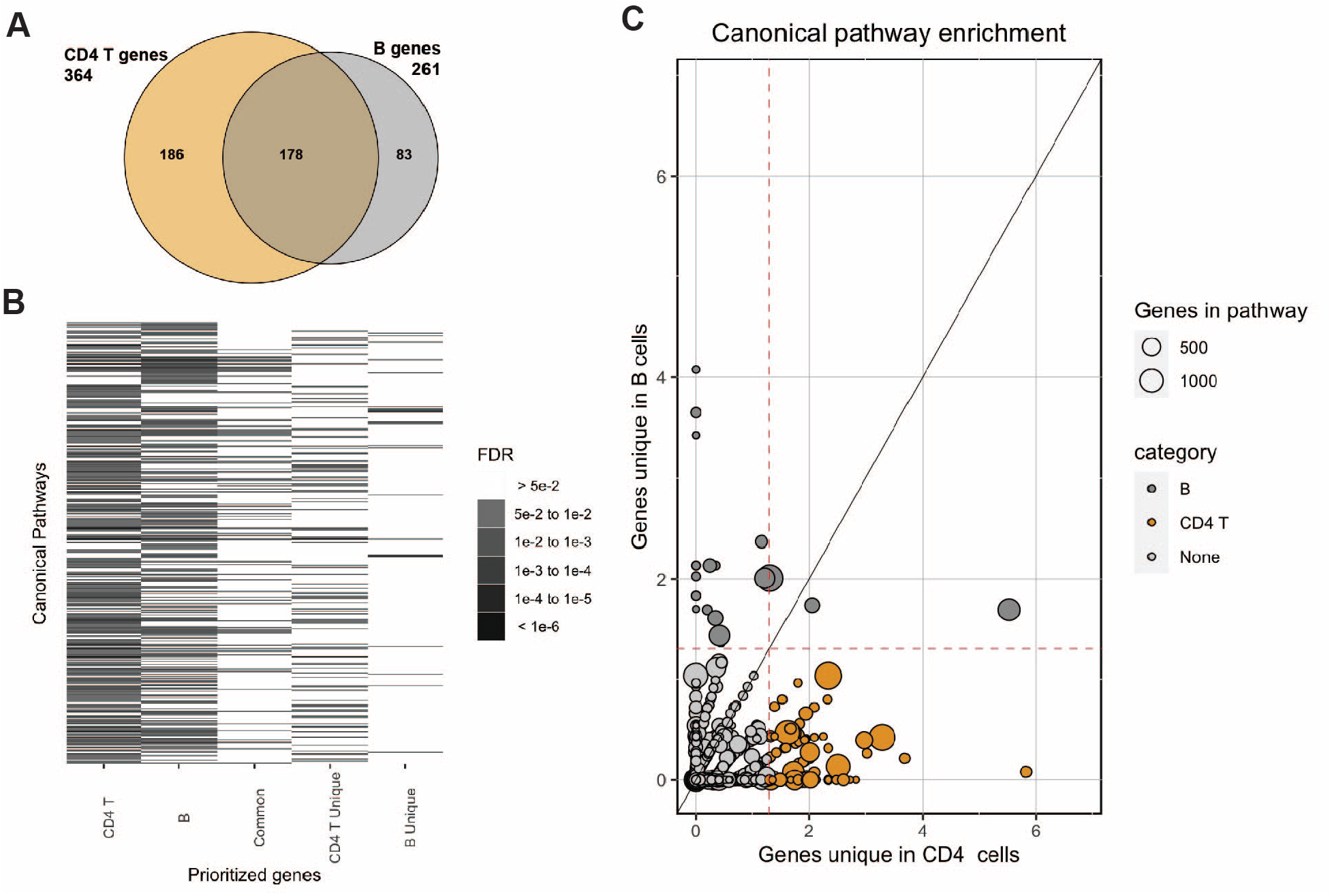
**A:** Venn diagram of the putative causal CD4 T and B cell genes. **B.** Heatmap of canonical pathway enrichment for the putative causal genes in CD4 T cells, B cells, common in CD4 T and B cells, unique in CD4 T cells, and unique in B cells. Only pathways with FDR<5% in at least one gene list are displayed (n=1950). The grayscale depicts level of statistical significance. **C.** Scatterplot of -log10(FDR) of canonical pathway enrichment for putative causal genes unique in CD4 T cells (X axis) vs. B cells (Y axis). The dashed red lines indicated FDR<5%. The size of the dots depicts the total number of genes in the respective pathway.

Next, we utilized the list of putative causal genes to identify enriched canonical pathways. Starting with CD4 T and B cell lists, we created additional lists for the common genes (shared between CD4 T and B cells), and finally genes unique to CD4 T cells and B cells. We observed widespread pathway enrichment for the putative causal genes of the CD4 T cells (n=294) and B cells (n=236) at FDR<5%. The common set of genes was enriched in 85 pathways (**Supplemental Table 25)**. The B unique gene list (n=83) was enriched in 22 canonical pathways, including lipoprotein and cholesterol pathways, the CD40 pathway, and JAK-STAT pathway (**Figure 7B-C**). The unique genes in CD4 T cells (n=186) were enriched in 99 pathways, including TCR pathways, various interleukin pathways, and MAPK/ERK signaling pathways (**Figure 7B-C**).

Pathway analyses utilize known biological connections for a given set of genes but many of underlying mechanisms could be still uncharacterized. Thus, we leveraged protein-protein interaction (PPI) data to test whether the respective putative causal gene lists exhibit a high degree of connectivity^38^. A similar percent of the mapped CD4 T cell and B cell prioritized genes were directly connected, 48.9% and 43.7% respectively (**Supplementary Table 26;** Figures S8**-12)**. Only the CD4 T and CD4 T unique gene lists demonstrated a higher degree of connectivity than expected (p-value<0.05), although all gene lists had communities of genes with high connectivity (p-value<0.05; **Supplementary Table 26**). These results are consistent with the pathway analyses, implying that the MS genetics are mediated by several different mechanisms in both cell types, some of which are shared and some of which are cell-type specific.

**Figure 8:**
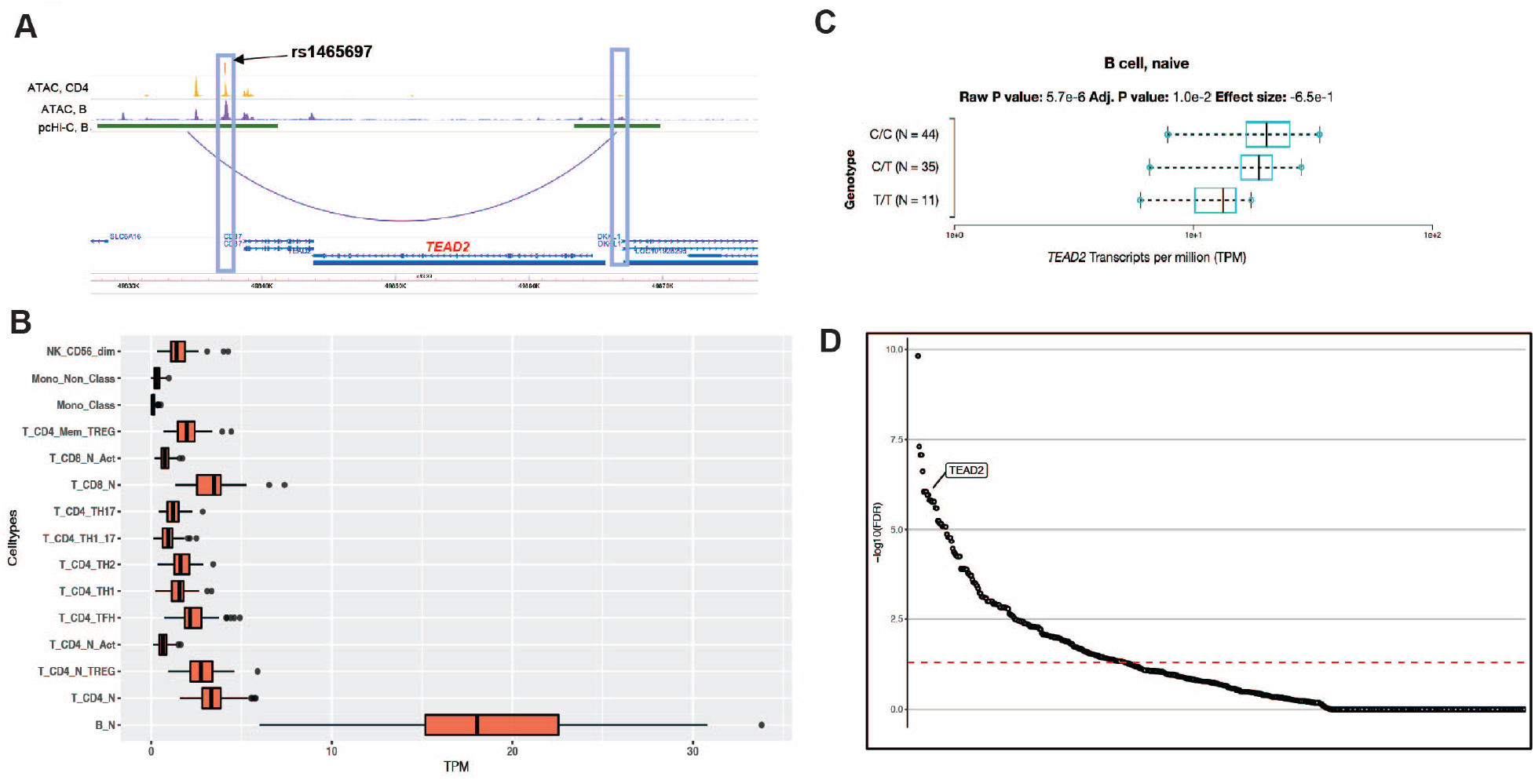
**A:** Visualization of *TEAD2* locus. Lead SNP rs1465697 (PICS of 15%) is depicted with a red line. The blue box on the left illustrates the overlap with the ATAC-seq peaks present in CD4 T (orange) and B cells (purple). The SNP and ATAC-seq peaks also overlap a PCHiC looping interaction with the promoter for the *TEAD2* gene (arc; the boundaries of the enhancer/promoter regions are indicated in green; the promoter of *TEAD2* is highlighted with the blue box on the right). **B**. Gene expression of *TEAD2* across immune cells available in the DICE database (https://dice-database.org/). X axis display transcripts per million (TPM). **C.** Cis-eQTL boxplot per genotype status of rs1465697 in naïve B cells in the DICE database (https://dice-database.org/). **D.** Transcription factor enrichment in the GTRD database for the putative causal genes that are common in CD4 T and B cells. Each dot represents one of 526 transcription factors. The Y axis indicates the -log10 of the FDR. The *TEAD2* enrichment is highlighted (p-value = 1.34x10^-8^, FDR = 8.81x10^-7^).

### A fine interplay between shared and cell-specific B and CD4 T cell putative causal genes

The results of the pathways and PPI analyses suggest that the MS genes common and unique to B and CD4 T cells do not act independently but rather have shared cellular mechanisms. We illustrate this by studying a locus on chromosome 19 (**Figure 8**) where the lead SNP rs1465697 (chr19:49837246 C>T) has a MS GWAS association p-value of 3.02x10^-18^. The lead SNP is the most highly prioritized variant by statistical fine-mapping out of 21 overall SNPs in the 95% credible set (PP=15% for rs1465697; next highest PP 9%). This SNP overlies an OCR present in all lymphoid lineages, including B cells, CD4 T cells, and CD8 T cells. We have previously suggested five putative causal genes for this locus: *DKKL1*, *CCDC155, CD37*, *TEAD2*, and *SLC6A16* ^8^. Using PCHiC data, this OCR forms a chromatin loop interaction with *TEAD2* and *DKKL1*, but not the other three genes. This chromatin loop interaction is observed in activated CD4 T cells, naïve B cells, total B cells, naïve CD8 T cells, and fetal thymus, but none of the other hematopoietic cell types ^37^. Furthermore, this SNP is an eQTL for *TEAD2* in B cells, but is not an eQTL for any other gene in this locus in any of the available hematopoietic cell types (**Figure 8**)^39^. Together, these lines of evidence support *TEAD2* as the causal gene at this locus.

Interestingly, *TEAD2* is a transcription factor with 1459 predicted regulated genes ^40^, including 38 putatively causal CD4 T cell genes and 25 B cell genes as nominated above (FDR < 1 %, FDR < 1%, respectively; **Figures S13-14**). The majority of these genes are common in both CD4 T and B cells (n=23; FDR < 1%; **Figure S15**). To identify genes whose expression is modulated by T*EAD2* in CD4 T and B cells, we examined *TEAD2* knock-down (KD) and over-expression (OE) in cancer cell lines (n=8, respectively) from the Library of Integrated Network-Based Cellular Signatures (LINCS) Program ^41^. Although these cell lines do not represent an ideal experimental model to study the effect of *TEAD2* in immune cells, they can still be used to understand mechanisms reflecting core cellular functions. Within each cell line we identified genes whose expression changed in the opposite direction (top or bottom 10%) in the KD versus OE models. These ranged from 7 to 24 for the CD4 T cell prioritized genes (**Supplementary Table 27; Figure S16**) and from 3 to 16 for the B cell genes (**Supplementary Table 28; Figure S17)**. Of these, 17 genes and 9 genes changed expression in at least 2 cell lines, respectively (**Supplementary Tables 27-28**). These data demonstrate that perturbation of *TEAD2*, a key immune cell transcription factor, results in indirect changes of putative MS causal genes in B and CD4 T cells.

## Discussion

In this paper, we integrated MS GWAS with chromatin accessibility data from a broad array of peripheral immune cells in order to identify putative causal cell types. Our analyses identified regulatory regions in B cells and CD4 T cells as each being independently enriched for MS genetics. Within the CD4 T cell and B cell populations, we further identified OCRs from T_h_17 cells and memory B cells as specifically driving their respective enrichments. Chromatin data from MS patients reiterated these findings and further suggested that immunomodulatory treatments alter the chromatin accessibility overlying MS-associated GWAS variants. Integration of PCHiC data led to prioritization of putative causal genes in B and CD4 T cells, identifying target genes that are both shared and specific to each of these cell populations. The putative causal genes implicate several known signaling pathways, mostly due to the cell-specific MS-associated genes, despite these representing a smaller percentage compared to genes shared between B and CD4 T cells. Finally, we illustrate that the B and CD4 T cells mechanisms are intertwined by describing how *TEAD2,* a putative causal gene shared between B and CD4 T cells, contributes to disease susceptibility directly and indirectly by targeting both shared and cell-specific MS genes.

Our study provides genetic evidence for the independent involvement of both CD4 T and B cells in the pathogenesis of MS ^1–3^ . It further supports the long-debated causal role of memory B cells in MS^24, 42, 43^, shown by the highly effective therapies targeting B cell receptors, most notably ocrelizumab^7^. Through our analyses, we also corroborate decades of research that have demonstrated a primary role of CD4 T cells in MS, including the importance of T_h_17 T cells ^24, 44, 45^. For both the B cell and T_h_17 T cell populations, we find that active enhancers and promoters drive the enrichment signal, consistent with activating roles of these cell types in MS as an autoimmune condition. The complex interplay between the B and CD4 cells in MS pathogenesis ^2, 3, 42^ is also reflected by our finding of some OCRs present in both cell types, while other OCRs are present only in specific cell populations.

Although other peripheral immune cell types have been shown to be involved in MS, including monocytes, CD8 T cells, and mDCs^24^, we did not identify independent enrichments for these cell types. One possible explanation for this apparent contradiction is that these cell types might work secondarily to memory B and T_h_17 cells, which are more directly under influence from MS GWAS-associated variants. Non-genetic effects, e.g. environment-specific response, could also explain their role in MS above and beyond any shared mechanisms with B and CD4 T cells. Further, context-specific studies, such as under various cell activation conditions, would be necessary to unravel any potential independent influence of these cells by MS genetic variants. Lastly, while our analyses do not identify an independent genome-wide enrichment for cell types other than B and CD4 T cells, GWAS variants may still act at individual loci in these other cell types.

One of the key challenges of GWAS is moving from genetic association to biological mechanisms^10, 13, 14^. This is driven by three main challenges. First, linkage disequilibrium, while highly advantageous to discovery of genetic associations, limits our ability to identify the causal variant. Second, as most GWAS variants are noncoding, identifying the gene(s) that are affected by the causal variant can be difficult. Third, the cell type(s) in which a given associated variant acts can be unclear. Our study demonstrates how we can use statistical fine-mapping to help solve the first challenge, though this is not without multiple caveats^46^. We integrated orthogonal datasets (ATAC-seq, PCHiC) to help delineate the likely causal genes and cell types and overcome the latter two challenges. We document how shared and cell-specific genes affect putative causal pathways. We further illustrate the complex interplay between shared and cell-specific putative causal MS genes by studying the *TEAD2* locus, a transcription factor recently implicated in immune regulation^47^. Together, our study generates important insights into the driver subpopulations of peripheral immune cells in MS, reinforcing how MS genetics act primarily through B and CD4 T cells. Our study also demonstrates the need for in-depth context-specific cellular data to carefully delineate the causal role of each immune cell subset in MS.

## Methods

### GWAS summary statistics

We utilized available MS GWAS summary statistics, which included data from 8,278,136 variants across 14,802 individuals with MS (cases) and 26,703 individuals without MS (controls) ^8^. For enrichment analyses, we included only the 6,773,531 variants that were analyzed in all 15 cohorts of the discovery stage meta-analysis; this resulted in 6,773,531 variants carried forward. We additionally utilized GWAS summary statistics from various neuropsychiatric disorders or autoimmune disorders: Alzheimer disease (AD)^26^, schizophrenia (SCZ)^27^, bipolar disorder (BPD)^28^, type 1 diabetes (T1D)^29^, Crohn’s disease (CD)^30^, ulcerative colitis (UC)^30^, systemic lupus erythematosus (SLE)^31^, rheumatoid arthritis (RA)^32^, and primary biliary cirrhosis (PBC)^33^. All GWAS data were converted to the “.sumstats” format as required by LDSC using the “munge.py” function in LDSC with default parameters.

### Epigenetic and eQTL datasets

#### Hematopoietic progenitor and terminal ATAC-seq data

ATAC-seq data for 16 different human hematopoietic progenitor and terminal populations was obtained from Corces et al.^18^ and Buenrostro et al.^19^. These ATAC-seq profiles were generated on bulk FACS-sorted cells from human peripheral blood or bone marrow cells. Alignment of the ATAC-seq data and peak-calling were performed as previously described^20^. To identify cell-type specific peaks for each cell type, we used ATAC-seq peaks for that cell type and removed any peaks that overlapped with a peak present in any one of the other 15 cell types. A single base pair overlap was considered to be overlapping.

#### Immune cell ATAC-seq data

We used publicly available immune cell ATAC-seq data (NCBI GEO GSE 118189) derived from flow-sorted peripheral blood cells ^21^. As each cell type had between 1-4 human donors, we merged the raw ATAC-seq data from the individual donors for a given cell type. We aligned ATAC-seq reads using bowtie2 version 2.2.1^48^ with default parameters and a maximum paired-end insert distance of 2000 base pairs. The bowtie2 index was constructed with the default parameters for the hg19 reference genome. We filtered out reads that mapped to the mitochondria and used samtools version 1.10^49^ to filter out reads with MAPQ < 30 and with the flags ‘-F 1804’ and ‘-f 2’. Additionally, duplicate reads were discarded using picard version 2.20.6 (http://broadinstitute.org.github.io/picard). Finally, chromatin accessibility peaks were identified with MACS2 version 2.1.1^50^ under default parameters and ‘-- nomodel --nolambda --keep-dup all --call-summits’.

#### ChIP-seq data

We downloaded available pre-processed ChIP-seq peak calls from ENCODE for B cells^34^. Where replicates were available, the bed files for the replicates were merged to create a composite set of peaks for each histone mark. Data for T_h_17 histone ChIP-seq were downloaded from the Roadmap Epigenomics Project^35^ (NCBI GEO GSM997225); we used pre-processed ChIP-seq peak calls generated in Amariuta et al^51^.

#### chromHMM

We used a 25 chromatin state model^36^, which are imputed based on 12 epigenetic marks from across 127 epigenomes generated as part of the Roadmap Epigenomics Project^35^. We used the chromatin states from B cells and Th17 CD4+ T cells. Chromatin states were downloaded from https://egg2.wustl.edu/roadmap/web_portal/chr_state_learning.html. We excluded the “Quiescent/Low” cell state, as it encompasses a large proportion of the genome, resulting in unstable estimates of heritability.

#### Verily ATAC-seq data

We utilized immune cell ATAC-seq data generated from Verily as part of the SysteMS collaboration with Brigham & Women’s Hospital (PIs: Dr. Chitnis and Dr. Weiner). Cell sorting: Frozen cryovials in liquid nitrogen were thawed in a 37℃ bead bath and centrifuged for 5 min at 600 x g, 4℃. The cell pellet was washed with 1 mL of FACS buffer, and the wash repeated. The cell pellet was resuspended in the residual volume with 2.5 uL of 0.33mg/mL S7 DNAse. 50 uL of staining cocktail was added for the respective flow cytometry panels to be analyzed (T cell, B cell, myeloid panel) and incubated for 25 min on ice and in the dark. Cells were washed in FACS buffer, resuspended in a final volume of 400 uL FACS buffer, and passed through a 35 µm cell strainer cap. Stained samples were sorted on a FACSAria Fusion (BD Biosciences, San Jose, CA). Using FACSDiva v8.0.1 software, the samples were gated first by forward and side scatter properties, then FSC-H vs FSC-A for singlet discrimination, and finally, with their respective markers for each cell type (**Supplementary Table 29**). For each cell type of interest, up to 500 cells were sorted into the tagmentation buffer.

ATAC-seq library preparation and sequencing: Cells were sorted directly into 20 uL of cold tagmentation buffer (10 uL TD, 2 uL 2% IGEPAL CA-630, 6 uL nuclease-free H2O, 2 uL TDE1 per sample), followed by incubation at 37℃ for 30 min with shaking at 500 RPM. Samples were stored at -20℃ until further processing. DNA was extracted with the QIAGEN MinElute PCR purification kit according to the manufacturer’s protocol, and samples were amplified with KAPA HiFi kits and Illumina Nextera indices. The amplified material was cleaned with the QIAGEN MinElute PCR purification kit and quantified using KAPA library quantification kits. Samples were normalized and pooled for sequencing on the NextSeq (Illumina).

Processing: Paired-end raw ATAC-seq reads were trimmed using NGmerge ^60^ using the default parameters. The reads were then aligned to GRCh38 using Bowtie2 version 2.3.5 ^49^. The resulting SAM files were converted and sorted into BAM format using samtools version 1.5 ^50^. We filtered out the reads with MAPQ<10 and reads that were aligned to mitochondria using samtools. In addition, duplicate reads were removed using picard version 2.20.6 (http://broadinstitute.org.github.io/picard). Finally, peaks were called using MACS2 version 2.2.5 ^51^ with default parameters and --keep-dup all --nomodel –nolambda. To obtain peaks for each cell type, we merged the peak files from all samples for that specific cell type using bedtools version 2.29 ^61^.

### Enrichment of GWAS results within ATAC-seq peaks

To calculate enrichments of the MS GWAS data within annotations (e.g., ATAC-seq or ChIP-seq peaks), we applied stratified LD SCore regression (LDSC)^16, 23^. LDSC was performed using LDSC v1.0.0 (https://github.com/bulik/LDSC), which was run on the discovery summary statistics from the MS GWAS discovery stage summary statistics^8^. The human MHC locus was excluded given its complex LD patterns as recommended by Finucane et al.^16^ To run LDSC, we used precomputed LD scores based on the European ancestry samples of the 1000 Genomes Project Phase 1^52^ which was restricted to HapMap3 SNPs^53^, and we generated partitioned LD scores for each set of annotations. To perform LDSC, we regressed the summary statistics (χ2) from a given GWAS on to annotation-specific LD scores, with baseline scores (original 53 annotation model), regression weights and allele frequencies based on 1000 Genome Project Phase 1 data as precomputed by software authors. We applied partitioned heritability analyses using LDSC under three different models:

1. To ask how much a given annotation contributes to trait heritability, we used a LDSC model that includes baseline annotations and an annotation of interest. The heritability enrichment of the annotation was defined as the proportion of SNP heritability in the category divided by the proportion of SNPs in that category; we report statistical significance of this enrichment as p-values.
2. When comparing multiple annotations (e.g., ATAC-seq peaks from different cell types), we ran a LDSC model that includes the baseline model and annotations from all cell types. In this scenario, we calculate for each annotation the coefficient *τ_c_* Which measures the contribution to SNP heritability for a given annotation to heritability in this overall model, stratified on other annotations in the model. Z-scores for the coefficient *τ_c_* were converted to a one-sided p-value, which we report as a measure of statistical significance.
3. We also performed LDSC on pairs of annotations, which we term pairwise stratified LDSC. In these models, we include the baseline model, an index annotation of interest, and a comparator annotation of interest. To run LDSC, we used a previously described extension of LDSC ^54^.

Throughout, all default LDSC parameters were used.

### Statistical fine-mapping

Statistical fine-mapping was performed using the marginal p-values from the replication (joint analysis) summary statistics from the MS GWAS ^8^. The 200 genome-wide significant loci at p<5x10^-8^ were used. LD was calculated between each lead variant and all variants with r^2^>0.2 and within a 2 Mb window based on the 1000 Genomes Phase 1 (European subset) reference panel^52^. PLINK v1.90b3.32 was used to perform LD calculations^55, 56^ with parameters of ‘--r2 --ld-window-kb 2000 --ld-window 999999 --ld-window-r2 0.2’.

We then applied PICS to each locus^15^. Briefly, PICS uses the lead association p-value and LD structure of the locus to calculate the most likely causal SNPs given the observed lead association signal. PICS probabilities represent the probability of a given SNP in a locus being the causal SNP. Default PICS parameters were used. From the PICS probabilities, we calculated 95% credible sets (CS). We defined the 95% CS as a set of variants such that the true causal variant has a 95% chance of being in the credible set. To calculate credible sets, for each locus, we ranked variants in descending order by their PICS probabilities. We then iteratively added variants to the credible set for that locus until the sum of their PICS probabilities was greater than or equal to 0.95. For CS inclusion, we also required the variant to have a PICS probability > 0.1.

### Identification of target genes

We leveraged promoter capture Hi-C (PCHiC) data from 17 hematopoietic cell populations to link genetic associations with genes that they may regulate ^37^. We filtered the PCHiC dataset for looping interactions with a CHiCAGO score > 5^57^. An overlap between a GWAS variant and a PCHiC looping interaction was considered if the GWAS variant overlapped any position in the non-promoter (“other end”) of the PCHiC interaction. For CD4^+^ T cells, we considered only GWAS SNPs that overlapped an ATAC-seq peak in bulk CD4^+^ T cells or any of the CD4^+^ T cell subsets (naïve effector CD4^+^ T cells, T_h_1, T_h_2, T_h_17, follicular T_h_, naïve T_regs_ and memory T_regs_), and which overlapped a PCHiC interaction in naïve CD4^+^ T cells (nCD4), total CD4^+^ T cells (tCD4), non-activated total CD4^+^ T cells (naCD4), or activated total CD4^+^ T cells (aCD4). For B cells, we considered only GWAS SNPs that overlapped an ATAC-seq peak in bulk B cells or any of the B cell subsets (naïve B cells, memory B cells, or plasmablasts), and which overlapped a PCHiC interaction in naïve B cells (nB) or total B cells (tB).

### Correction for multiple hypothesis testing

Throughout our manuscript, we use Bonferroni corrections when testing multiple hypotheses. To generate a Bonferroni-corrected p-value threshold, we used a traditional p-value threshold of 0.05 divided by the number of tests being performed in a given analysis. We note that as many of the tests are correlated (since the underlying annotations are often highly correlated with each other), the effective number of independent tests being performed is fewer than the number of tests actually performed. As such, our analyses are overly conservative. We decided on this approach of using Bonferroni corrections as opposed to false discovery rate (FDR) approaches, as the number of tests being performed is often small, leading to unstable estimates of FDR.

### Gene set enrichment analyses

We performed pathway analyses utilizing the canonical pathways (CP) of the Molecular Signatures Database (MSigDB v7.2), as it is available from the Gene Set Enrichment Analysis website (http://software.broadinstitute.org/gsea/msigdb). We ran the Canonical Pathways, Biocarta, KEGG, and Reactome gene sets categories together in the same model. We estimated statistical significance using the hypergenometric distribution and applied false discovery correction, as previously described^8^. The same model was applied for the enrichment of prioritized gene sets with Gene Transcription Regulation Database (GTRD) transcription factor targets gene sets^40^. Significant enrichment level was set to a false discovery rate < 5%.

### Protein-protein interaction networks

We utilized GeNets (https://apps.broadinstitute.org/genets)^38^ to leverage known protein-protein interactions (PPI) of our prioritized gene sets. GeNets uses a random forest classified, trained in PPI data with 18 parameters that capture information about centrality and clustering. It creates communities of genes, sets of genes (nodes) that are connected to each other more than genes outside this community. Furthermore, it uses the random forest classifier and the connectivity to the tested gene set to propose candidate genes. For each described network the p-value is estimated by testing whether the number of observed edges divided by the numbers of possible edges using permutations. We ran GeNets via the web interface with the GeNets Metanetwork v1.0 and utilizing the InWeb model (“Override network the analysis model was trained on” option).

### TEAD2 knockdown and over-expression in LINCS cell lines

Robust z-scores, “level 5 data”, from knock-down (KD) or over-expression (OE) of *TEAD2* in cancer cell lines^41^ were downloaded from clue.io (https://clue.io/command?q=/sig%20%22TEAD2%22). The robust z scores represent differential expression for each genetic perturbagen, adjusted for the gene expression of all other perturbagens on the same physical plate. For knockdown and over-expression experiments the differential expression comparator were samples using a vector control, which are negative genetic controls that either lack a gene-specific sequence or target a non-human gene (like GFP).

## Data availability

MS GWAS summary statistics are available via request to the IMSGC (https://imsgc.net/). LD score regression software and reference panels were obtained from software developers (https://github.com/bulik/LDSC). Processed GWAS summary statistics for diseases other than MS were obtained from https://alkesgroup.broadinstitute.org/LDSCORE/independent_sumstats/. 1000 Genomes Phase 1 reference panel was obtained from ftp://ftp.1000genomes.ebi.ac.uk/vol1/ftp/. Hematopoietic ATAC-seq data for Buenrostro et al was obtained from NCBI GEO (accession GSE74912). Hematopoietic ATAC-seq data from Calderon et al was obtained from NCBI GEO (accession GSE118189). ChIP-seq data from Roadmap were obtained from NCBI GEO (accession GSM997225). chromHMM chromatin state partitions for ENCODE were obtained from https://egg2.wustl.edu/roadmap/web_portal/chr_state_learning.html. PCHiC data were downloaded from Data S1 in the referenced manuscript (https://www.sciencedirect.com/science/article/pii/S0092867416313228). DICE eQTL data were obtained from https://dice-database.org/.

The Verily Life Sciences ATAC-seq from the SysteMS data can be requested by Charlie Kim.

The LINCS KD and OE TEAD2 data can be accessed through the Broad Connectivity Map portal at clue.io: https://clue.io/command?q=/sig%20%22TEAD2%22.

## Acknowledgements

NAP was supported in part by National Multiple Sclerosis Society (grants JF-1808-32223 and RG-1707-28657). This work was supported in part by the Water Cove Charitable Foundation. We thank Jacob Ulirsch for technical assistance with the use of LD score regression.

## Author contributions

MHG and NAP had the original idea and supervised the project. JB, CC, XD, PK, CCK, TO, TS, and DZS generated data. HLW and TC provided samples. MHG, PS, BAL, HL, and NAP performed analyses. MHG and NAP wrote a first draft of the paper. All authors reviewed results, contributed to writing and final approval of the paper.

## Competing interests

JB, CC, XD, PK, CCK, TO, TS, and DZS employment in Verily Life Sciences at time of study.

## Computer code

Code used in this paper can be accessed in bitbucket: https://bitbucket.org/patslab/pis_ms_enrichment/

## Supplemental Figure Legends

**Figure S1:**
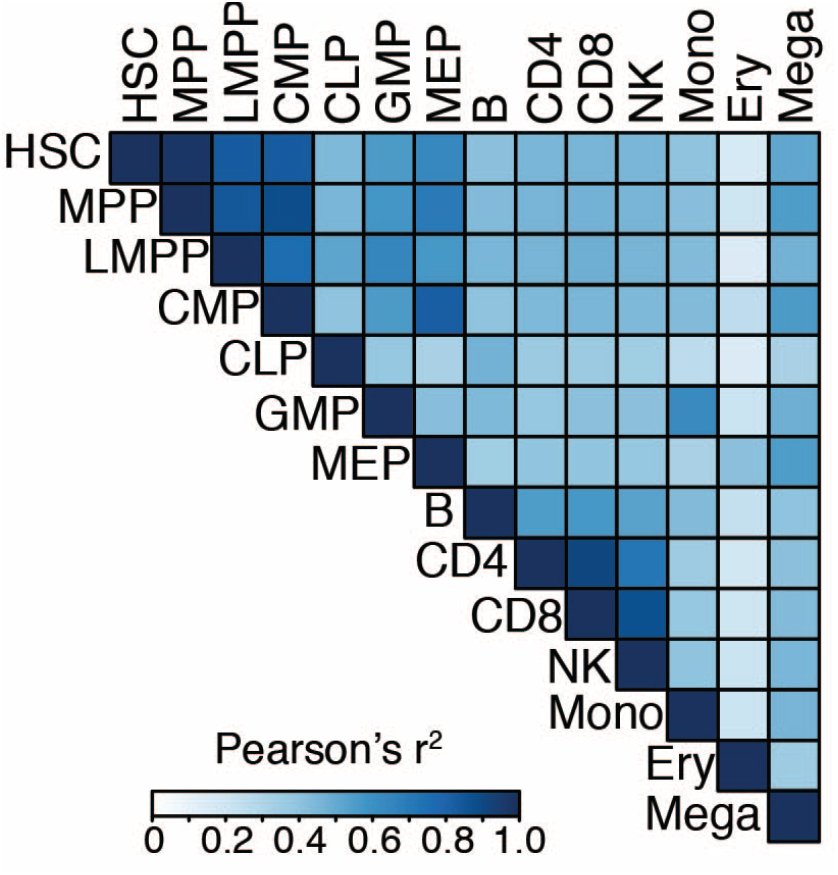
Correlation (Pearson’s r^2^) in ATAC-seq profiles across hematopoietic cell types.

**Figure S2:**
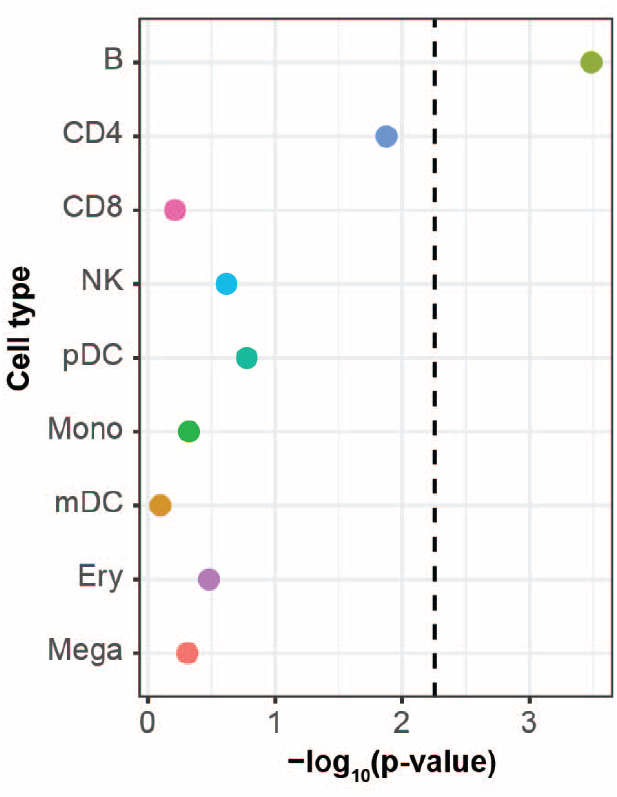
LDSC enrichments for MS GWAS in cell-type specific ATAC-seq peaks in each of the nine mature hematopoietic cell type. Y-axis shows –log_10_(p-value) of the LDSC heritability enrichment.

**Figure S3:**
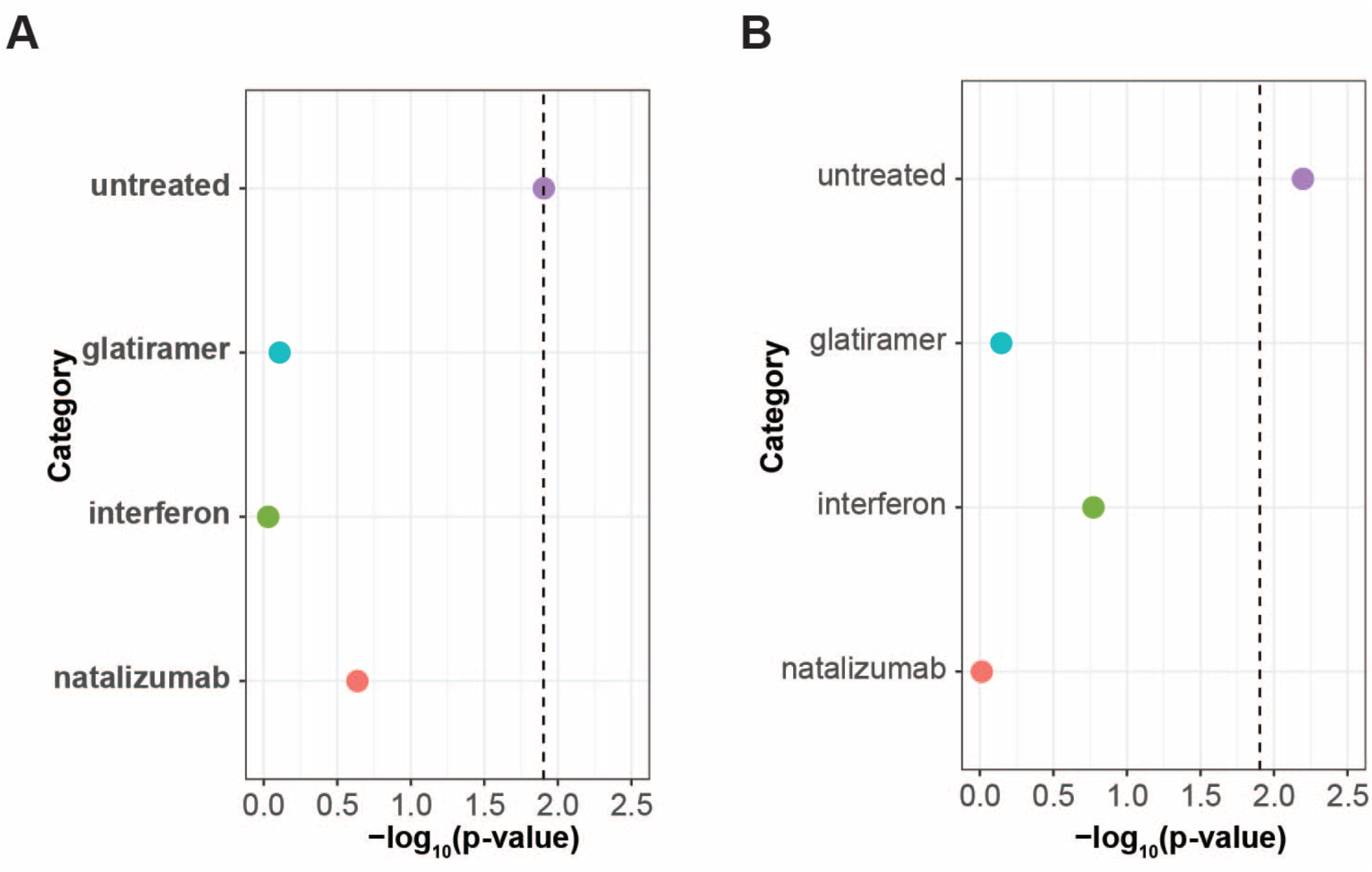
**A:** LDSC enrichment results for MS GWAS enrichment in T4cm OCRs from treated and MS treated patients in a joint model. **B.** Stratified LDSC enrichment results for MS GWAS enrichment in cMBc OCRs from treated and MS treated patients in a joint model. Heights of the circles reflect stratified LDSC coefficient p-values. Sizes of the circles are proportional to the enrichment p-values for that given cell type, with larger circles reflecting more significant p-values.

**Figure S4:**
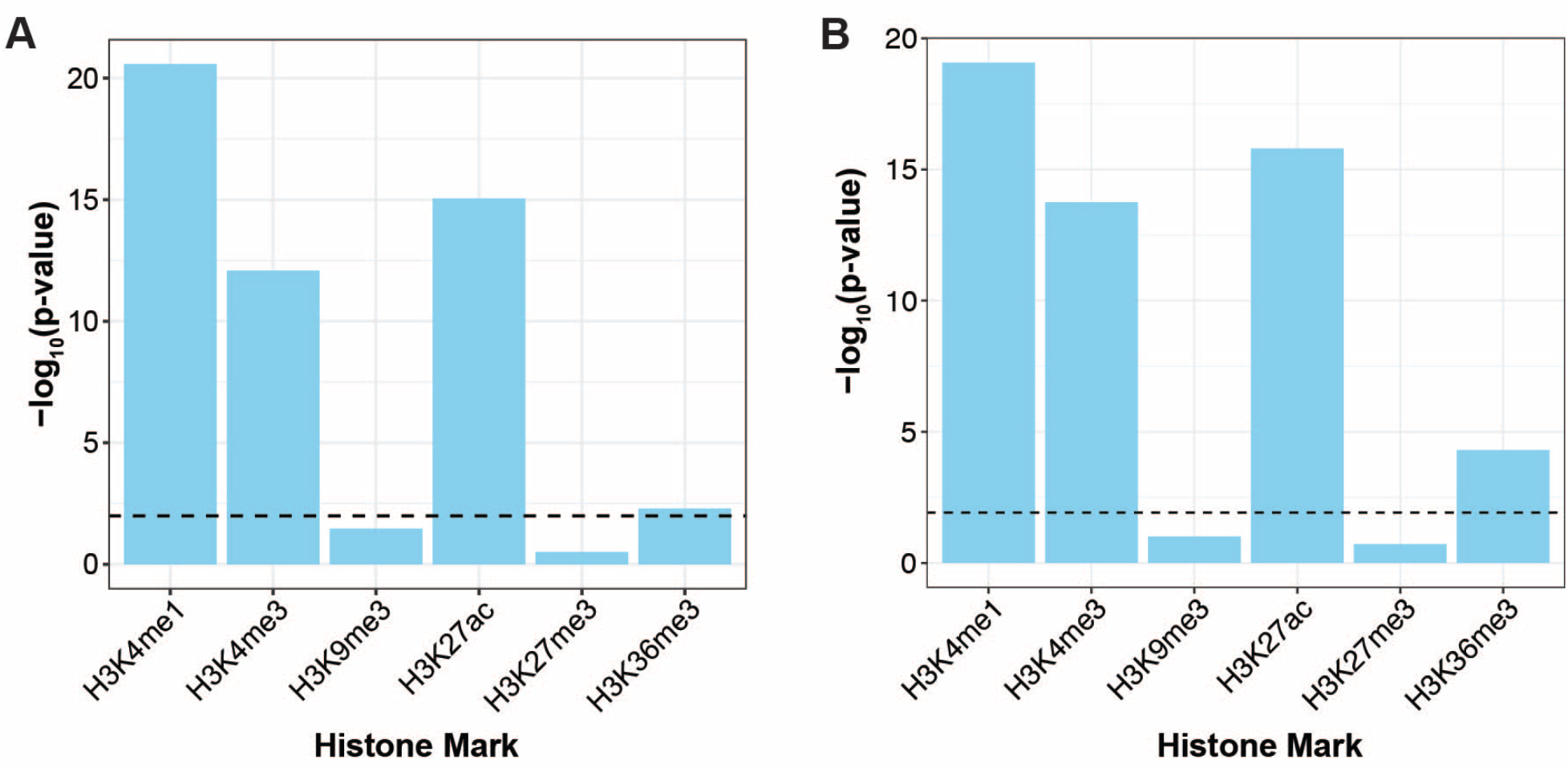
**A:** LDSC enrichment p-values for MS GWAS data in CD4^+^ T cell ChIP-seq peaks of various histone markers. Y-axis shown as –log10(p-value). **B:** Same as Figure S3A, except performed for B cell ChIP-seq histone markers.

**Figure S5:**
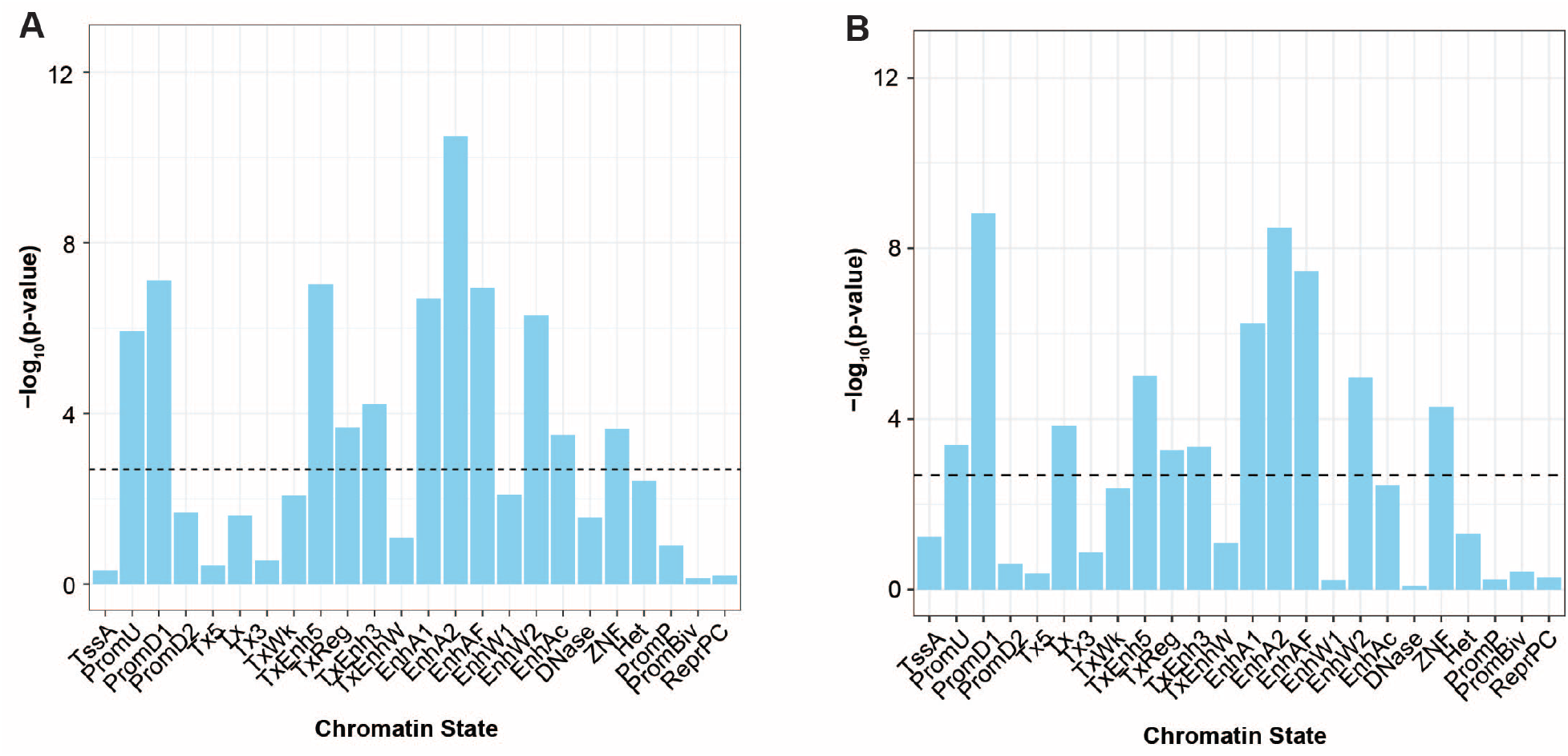
**A:** LDSC enrichment p-values for chromHMM chromatin states in CD4+ T cells from ENCODE. **B:** Same as Figure S5A, except for B cells from ENCODE. TssA: Active TSS; PromU: Promoter Upstream TSS; PromD1: Promoter Downstream TSS 1; PromD2: Promoter Downstream TSS 2; Tx5: Transcribed - 5’ preferential; Tx: Strong transcription; Tx3: Transcribed - 3’ preferential; TxWk: Weak transcription; TxReg: Transcribed & regulatory (Prom/Enh); TxEnh5: Transcribed 5’ preferential and Enh; TxEnh3: Transcribed 3’ preferential and Enh; TxEnhW: Transcribed and Weak Enhancer; EnhA1: Active Enhancer 1; EnhA2: Active Enhancer 2; EnhAF: Active Enhancer Flank; EnhW1: Weak Enhancer 1; EnhW2: Weak Enhancer 2; EnhAc: Primary H3K27ac possible Enhancer; DNase: Primary DNase; ZNF: ZNF genes & repeats; Het: Heterochromatin; PromP: Poised Promoter; PromBiv: Bivalent Promoter; ReprPC: Repressed Polycomb.

**Figure S6:**
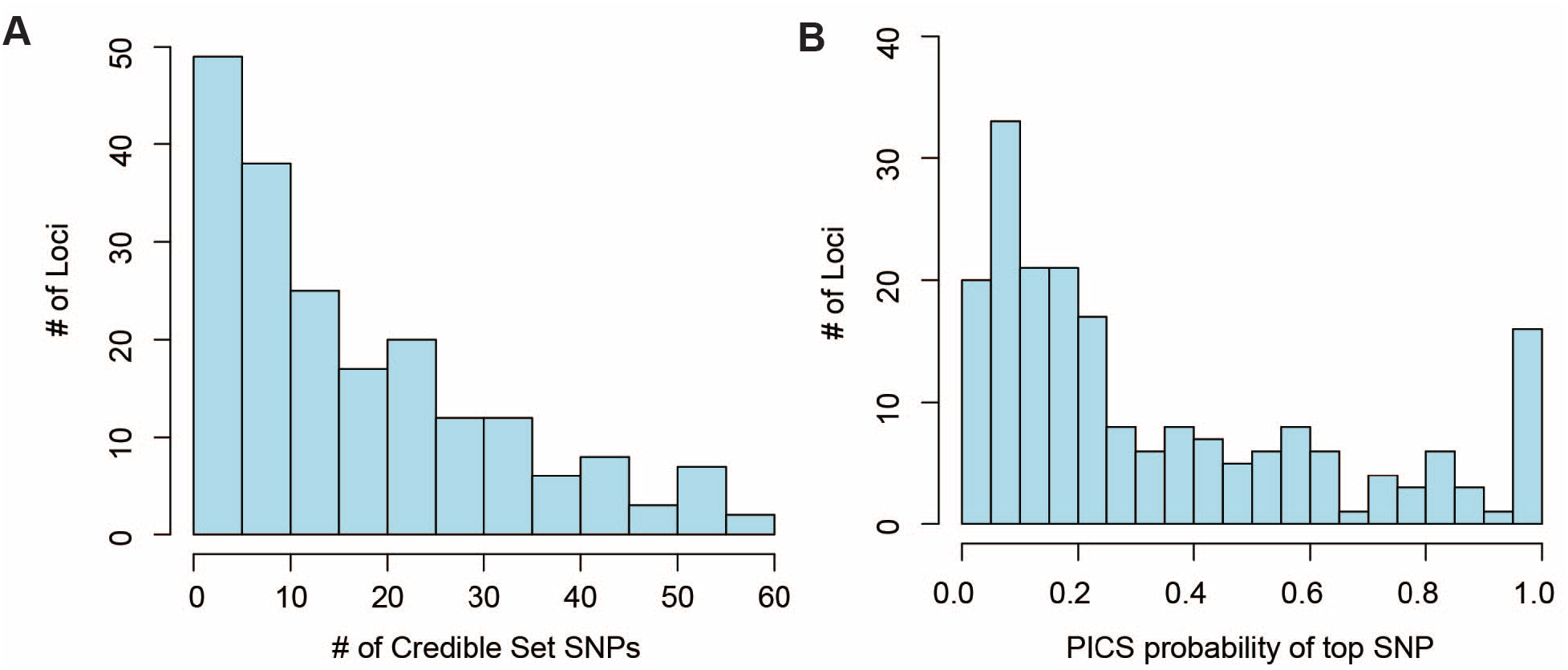
**A:** Histogram of the number of credible set (CS) variants across loci. **B:** Histogram of the PICS probability of the top variant in each locus.

**Figure S7:**
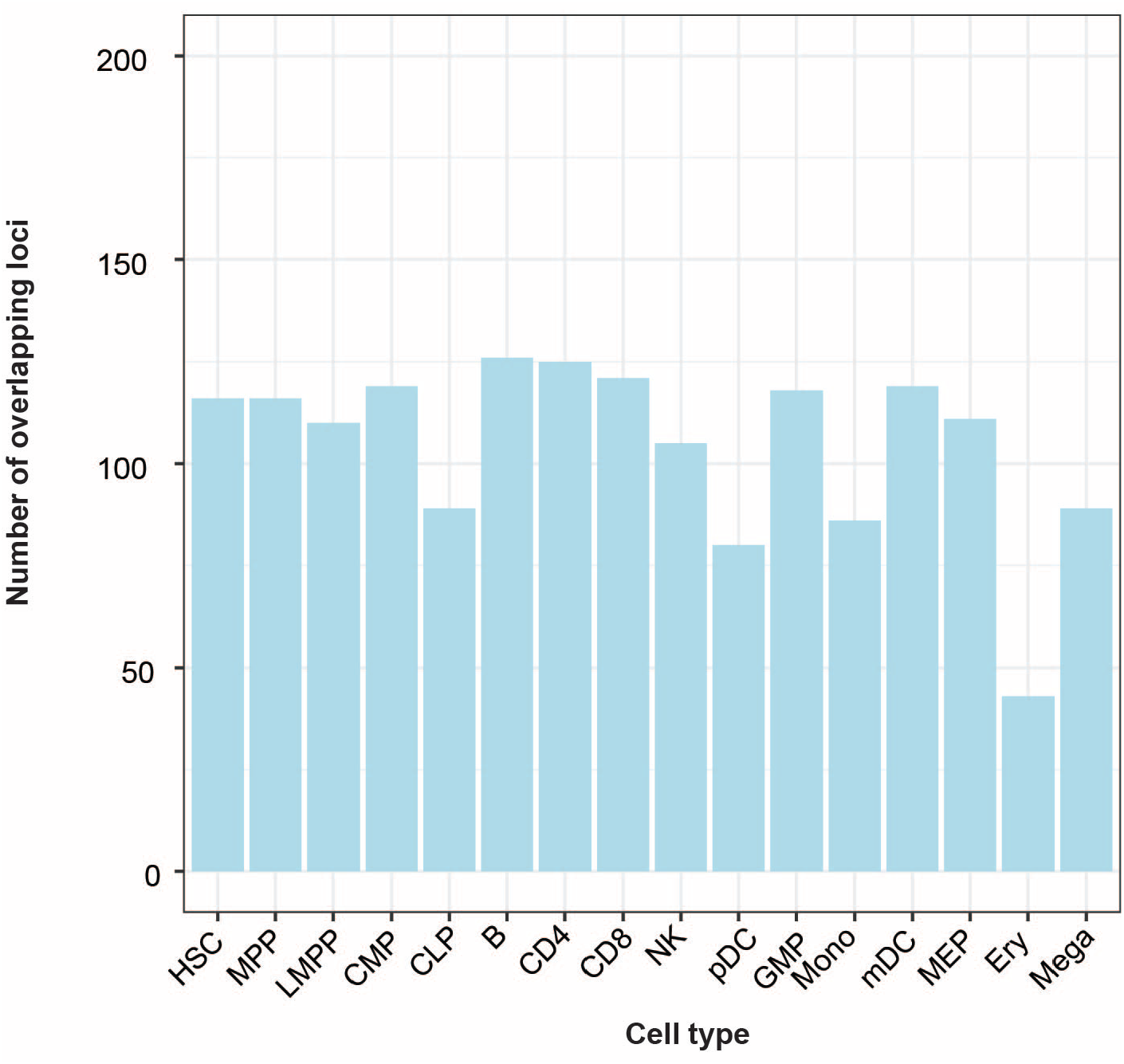
Number of MS GWAS loci (out of 200) with at least one CS SNP overlapping an ATAC-seq peak in the listed cell types.

**Figure S8:**
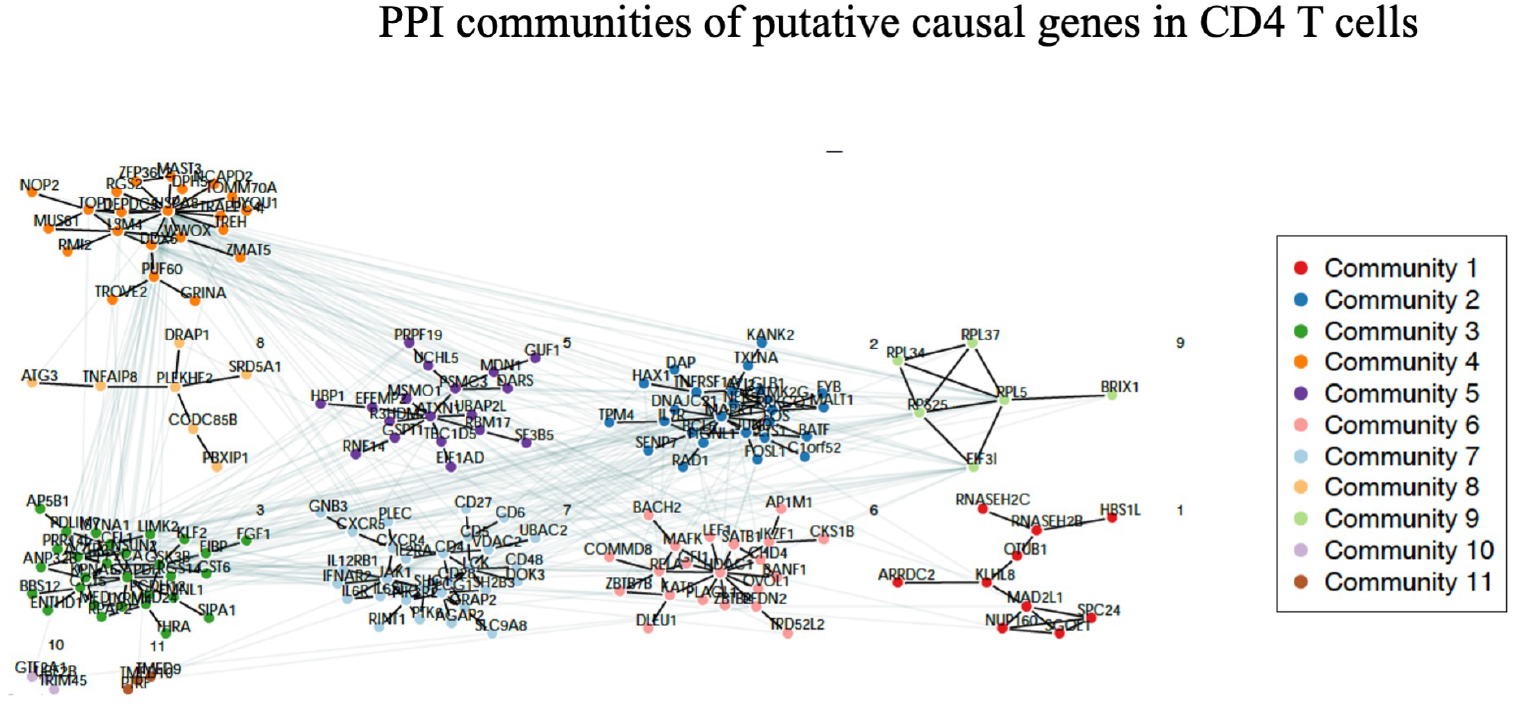
Protein-protein interaction communities of putative causal genes in CD4 T cells.

**Figure S9:**
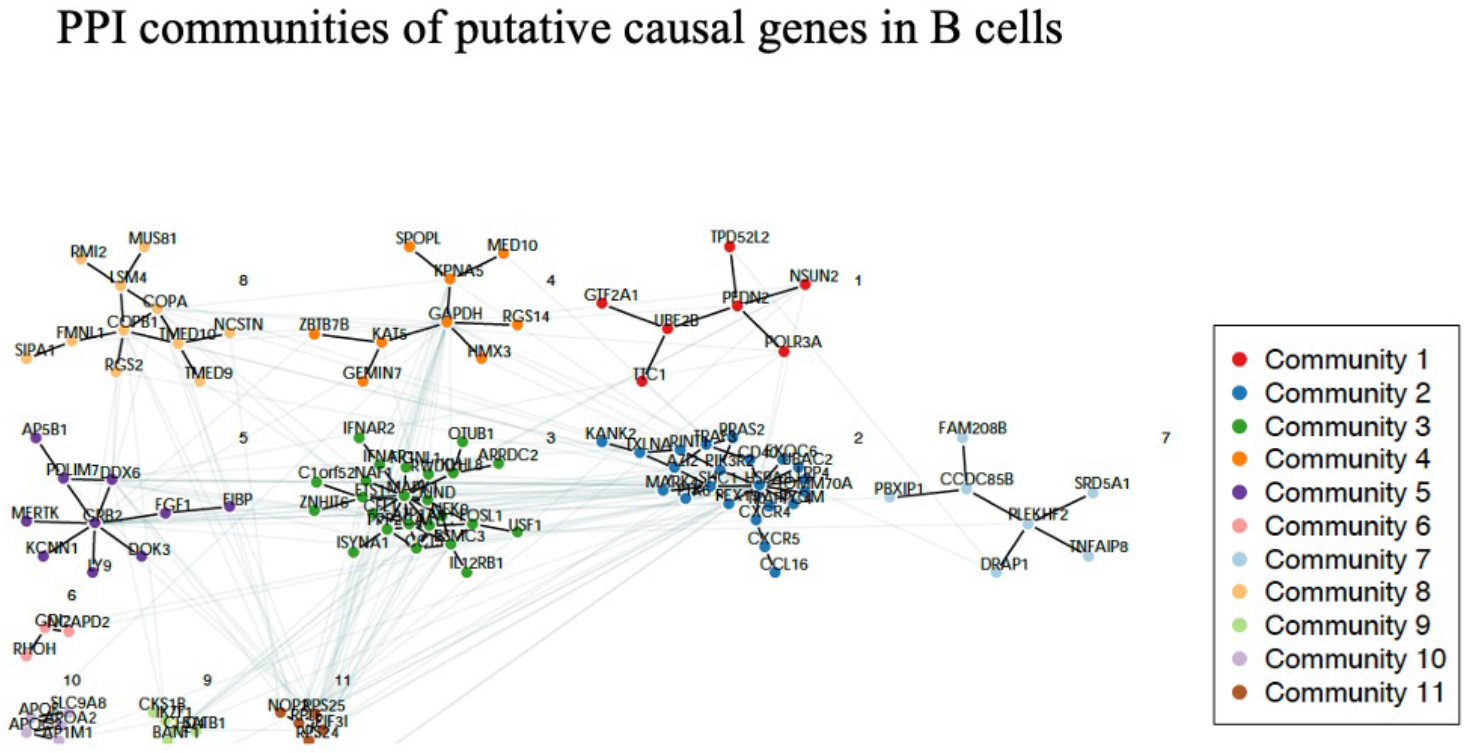
Protein-protein interaction communities of putative causal genes in B cells.

**Figure S10:**
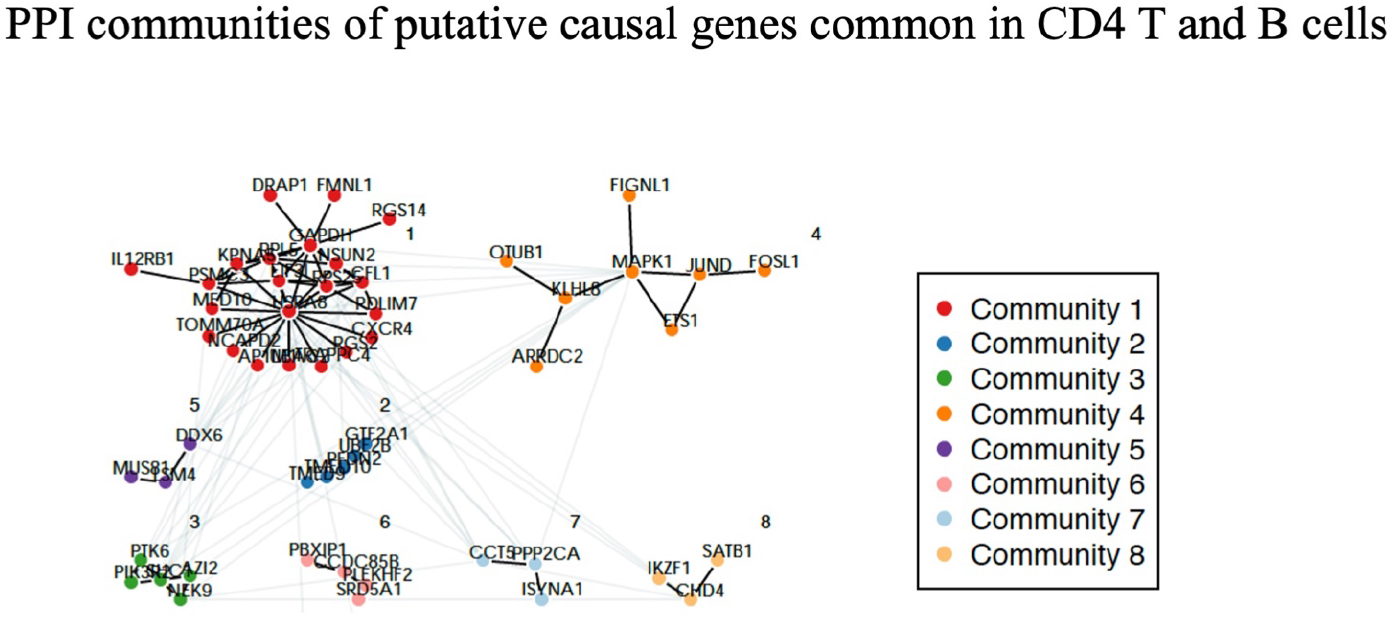
Protein-protein interaction communities of putative causal genes shared in CD4 T and B cells.

**Figure S11:**
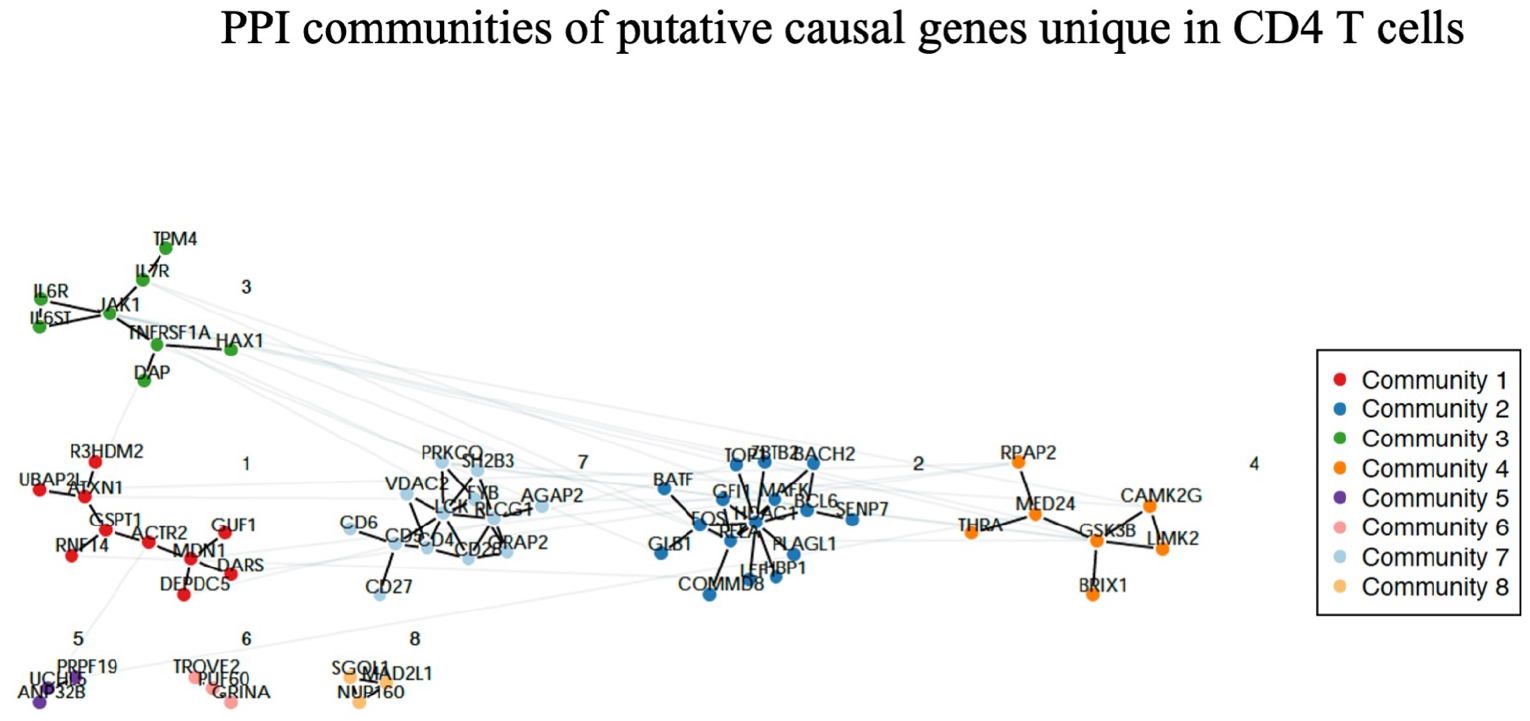
Protein-protein interaction communities of putative causal genes unique in CD4 T cells.

**Figure S12:**
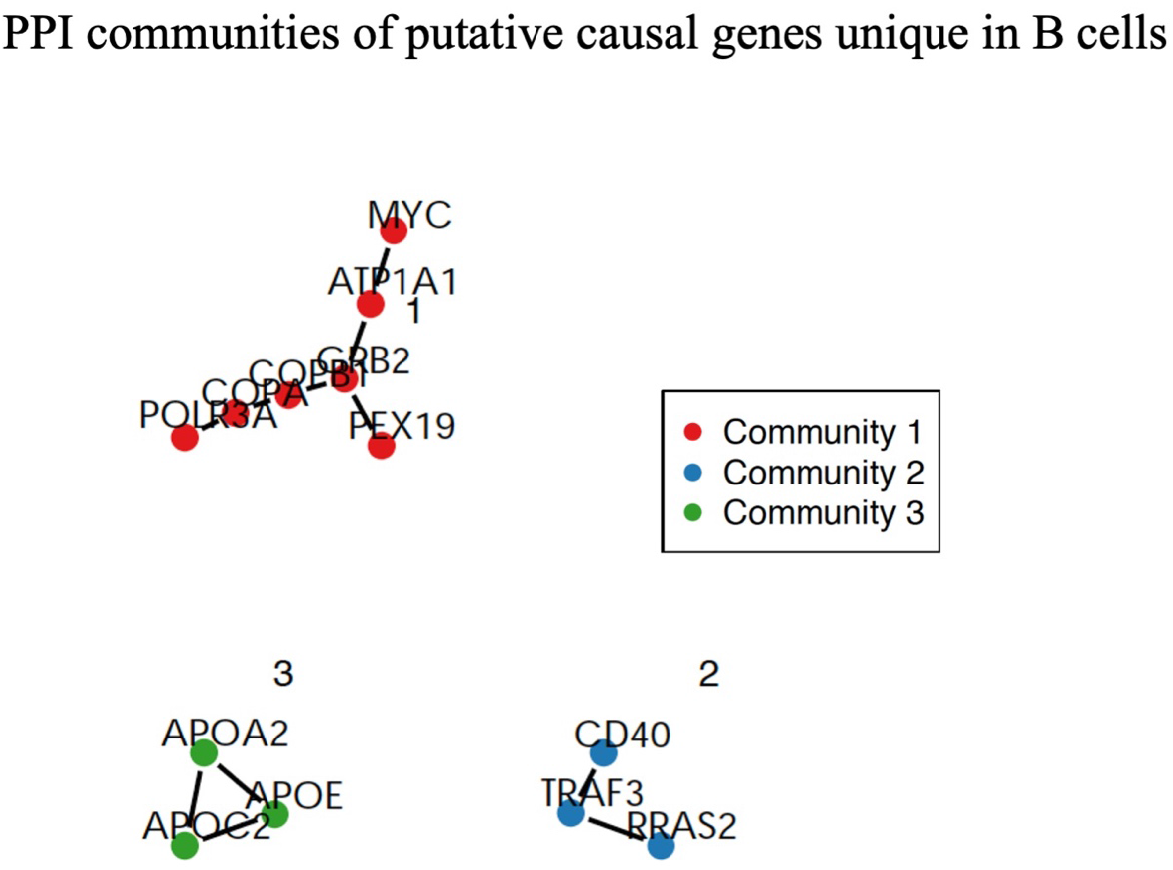
Protein-protein interaction communities of putative causal genes unique in B cells.

**Figure S13:**
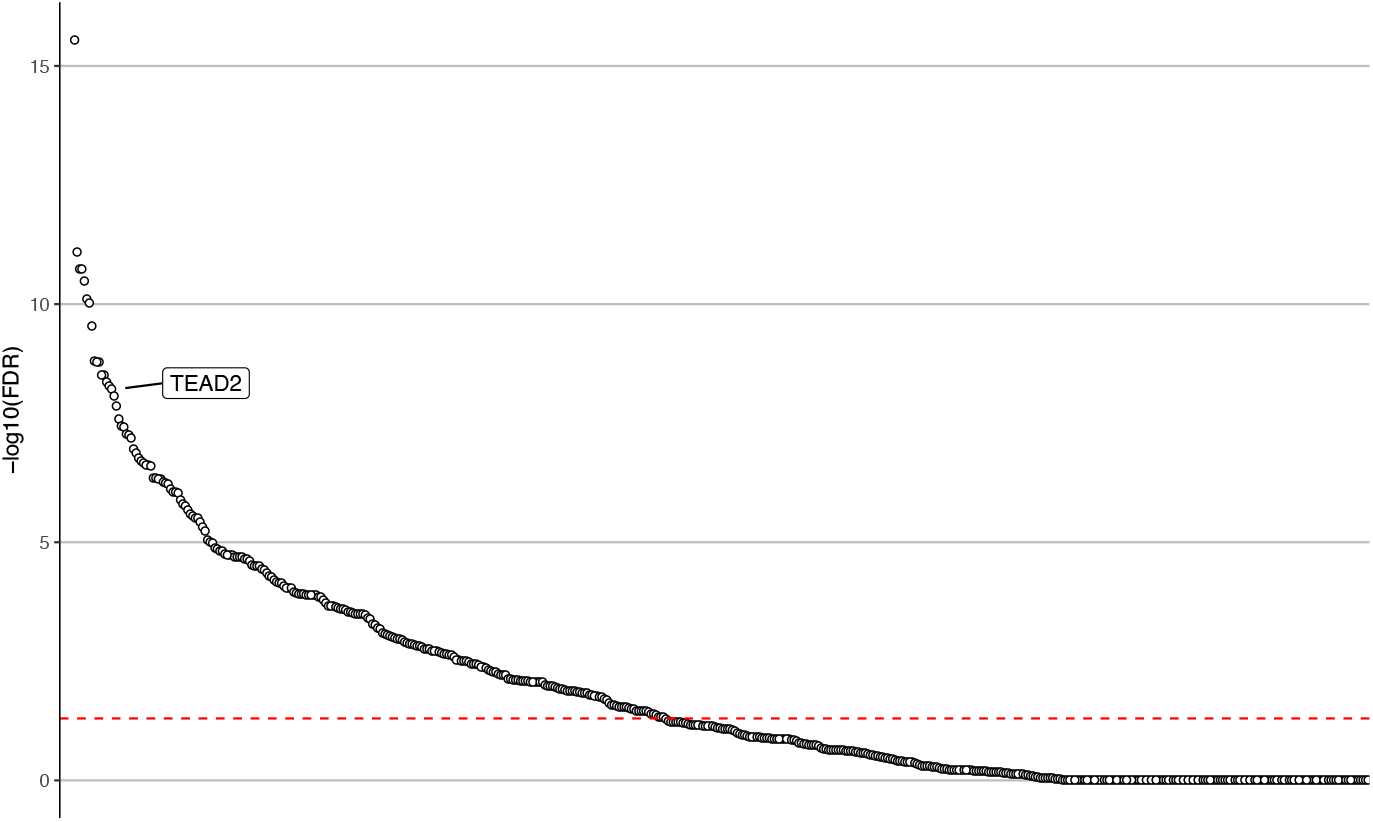
Enrichment of CD4 T cell putative causal genes in GTRD database. Each dot represents one transcription factor. The Y axis displays -log10 of false discovery rate (FDR). The dashed red line indicates the threshold of 1% FDR. The enrichment for TEAD2 predicted target genes is labeled.

**Figure S14:**
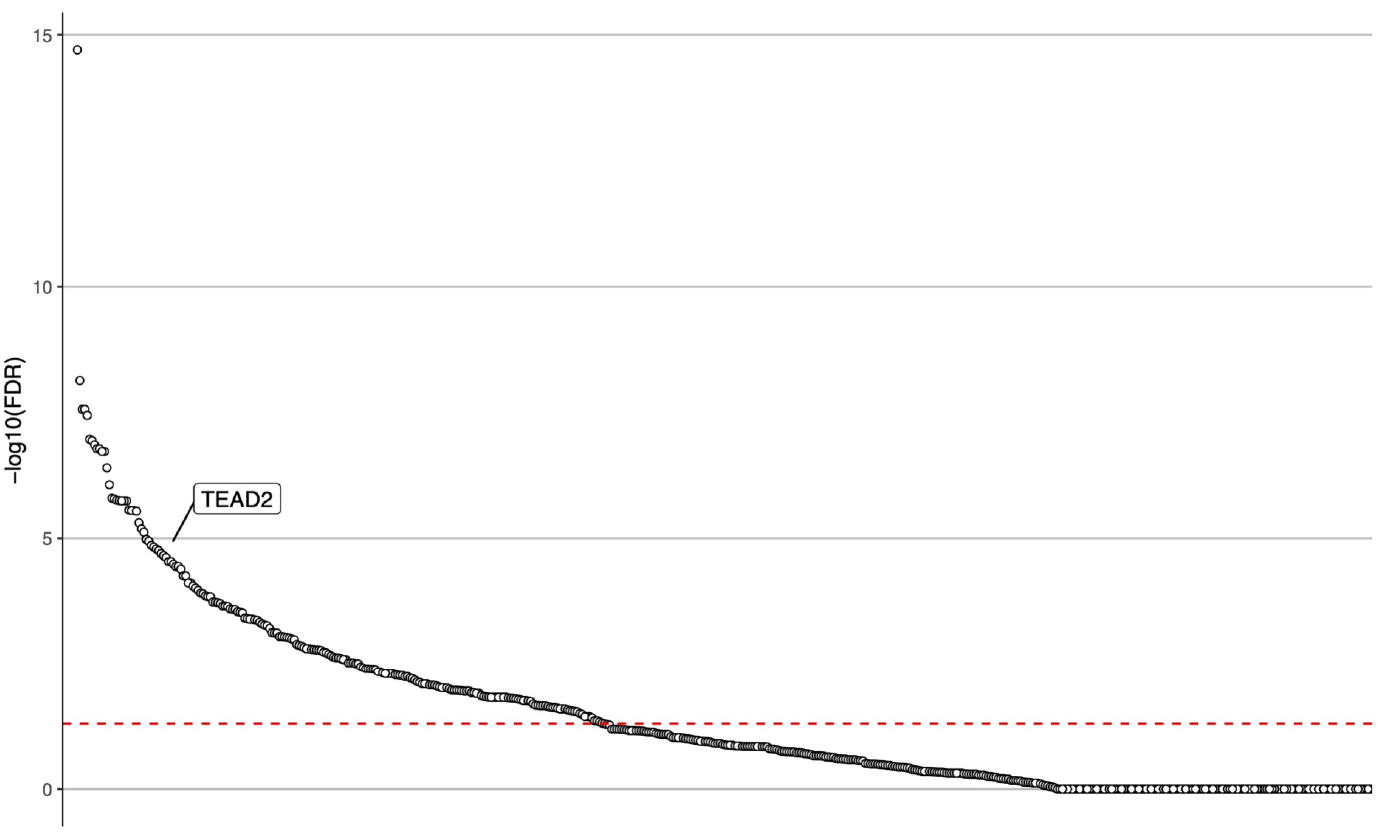
Enrichment of B cell putative causal genes in GTRD database. Each dot represents one transcription factor. The Y axis displays -log10 of false discovery rate (FDR). The dashed red line indicates the threshold of 1% FDR. The enrichment for TEAD2 predicted target genes is labeled.

**Figure S15:**
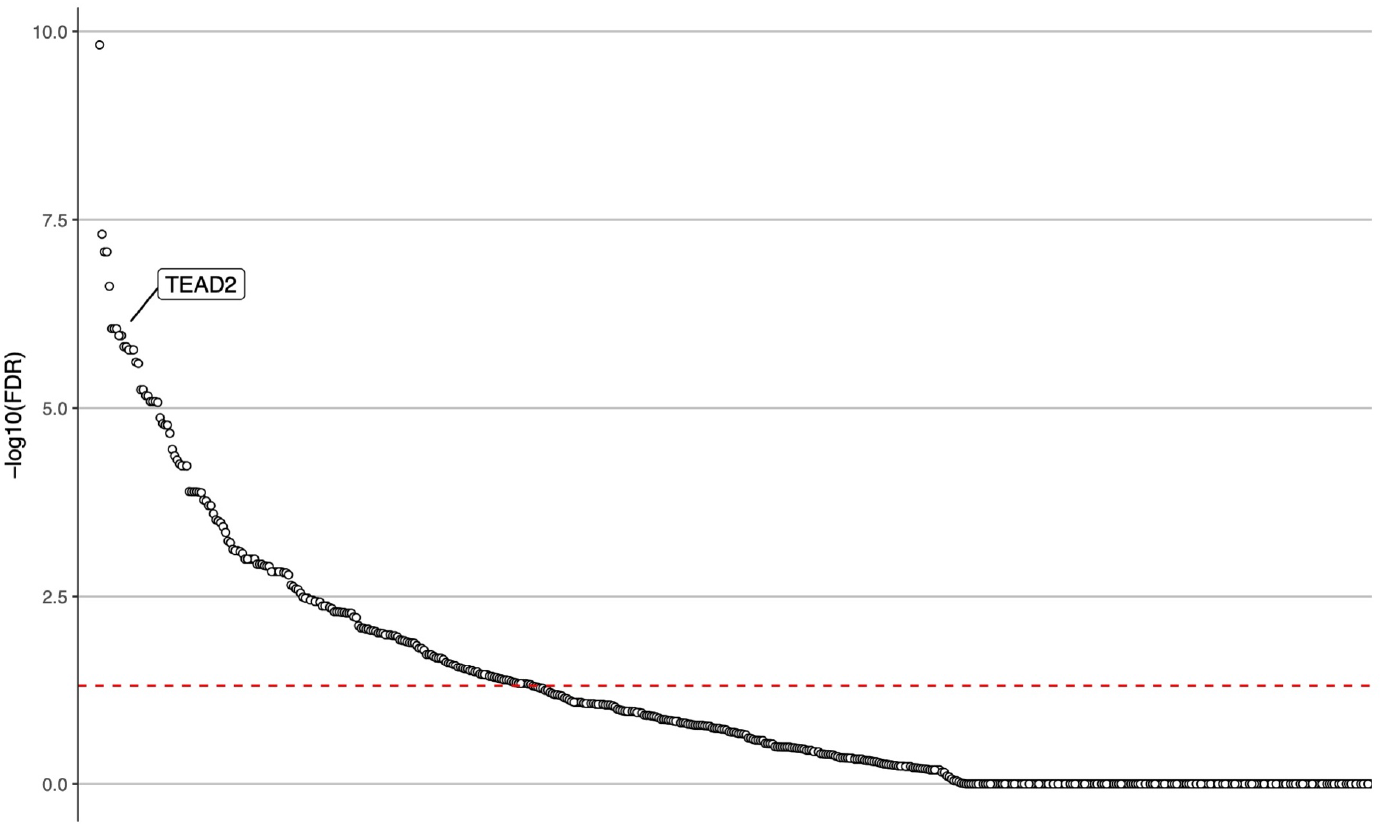
Enrichment of shared between CD4 T and B cells putative causal genes in GTRD database. Each dot represents one transcription factor. The Y axis displays -log10 of false discovery rate (FDR). The dashed red line indicates the threshold of 1% FDR. The enrichment for TEAD2 predicted target genes is labeled.

**Figure S16:**
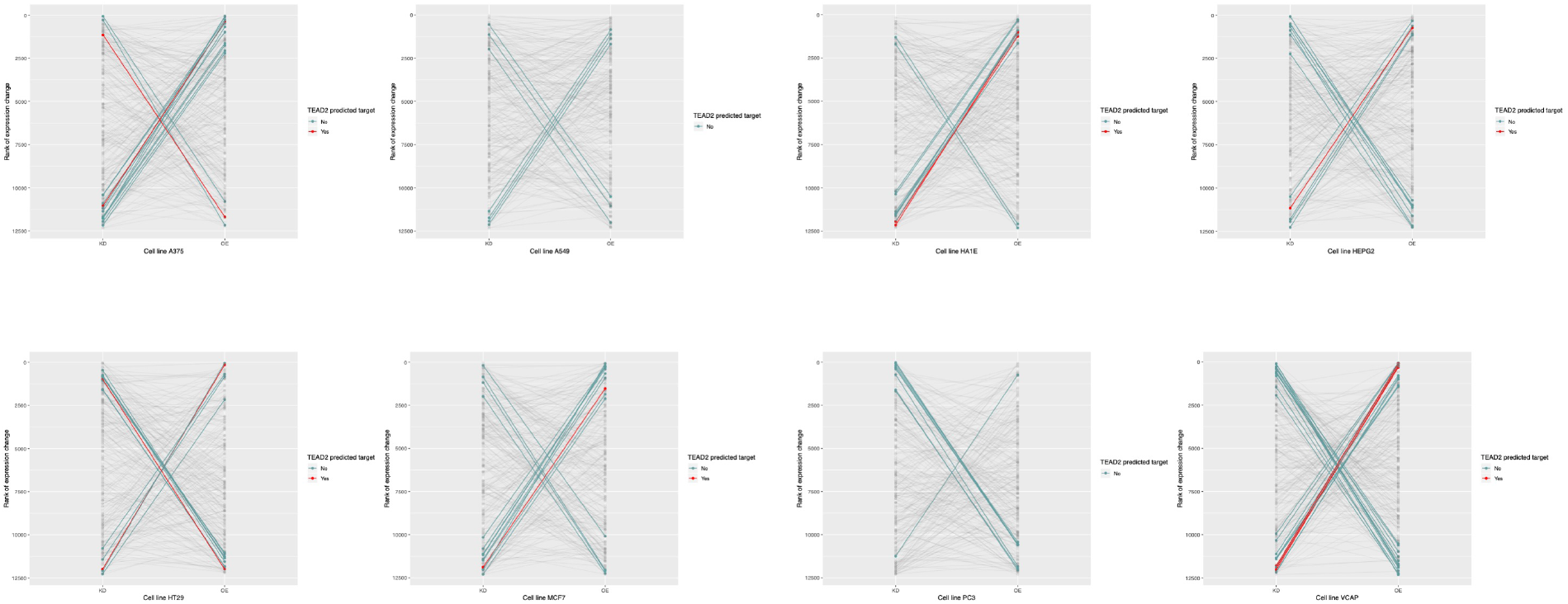
Change of gene expression of CD4 T cell putative causal genes in knock-down (KD) and over-expression (OE) models in cancer cell lines. Eight cancer cell lines are displayed: A375, A549, HA1E, HEPG2, HT29, MCF7, PC3, and VCAP. Putative causal genes are represented with lines connecting the ranked KD gene expression data (left column) with the ranked OE gene expression data (right column). Genes that are in the extreme 10% in opposite directions are indicated with green solid lines or red solid lines if these are also a predicted gene target for TEAD2. The light grey lines display all over putative causal genes.

**Figure S17:**
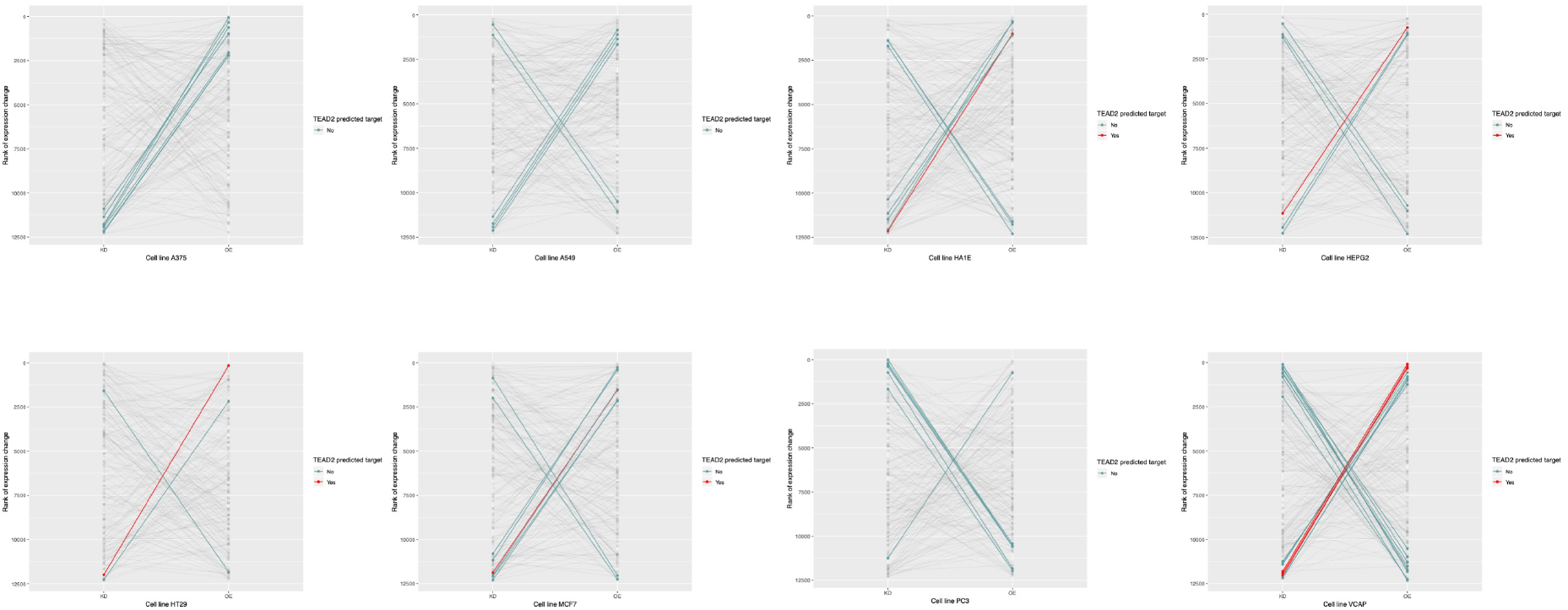
Change of gene expression of B cell putative causal genes in knock-down (KD) and over-expression (OE) models in cancer cell lines. Eight cancer cell lines are displayed: A375, A549, HA1E, HEPG2, HT29, MCF7, PC3, and VCAP. Putative causal genes are represented with lines connecting the ranked KD gene expression data (left column) with the ranked OE gene expression data (right column). Genes that are in the extreme 10% in opposite directions are indicated with green solid lines or red solid lines if these are also a predicted gene target for TEAD2. The light grey lines display all over putative causal genes.

## Supplementary Tables

**Supplementary Table 1: MS GWAS enrichment in 16 hematopoietic cell types.** Results from LDSC from a model including annotation of interest and set of baseline annotations. Column 2 shows the Proportion of SNPs in OCRs of that cell type. Column 3 shows the SNP heritability (h^2^_g_) for the annotation. Column 4 shows the standard error for h^2^g. Column 5 shows enrichment of SNP heritability, defined as proportion of SNP heritability in the annotation divided by the proportion of SNPs in that annotation. Column 6 shows the standard error of the enrichment of SNP heritability. Column 7 shows the p-value of the enrichment of SNP heritability.

**Supplementary Table 2: MS GWAS enrichment of hematopoietic cell types in joint model:**

Results from LDSC from a joint model including annotations from all cell types and set of baseline annotations. Column 1 shows the coefficient *τ_c_* which measures the contribution for a given annotation to heritability in this overall model, stratified on other annotations in the model. Column 2 shows the standard error for the coefficient *τ_c_*. Column 3 shows the z-score for coefficient *τ_c_* and column 4 shows the p-value for coefficient *τ_c_*.

**Supplementary Table 3. MS GWAS pairwise enrichment in 16 hematopoietic cell types.** Index cell types are shown on the left. The comparator cell types are shown on top. Coefficient p-values are shown for the index cell type in a model that includes the index cell type, comparator cell type, and the set of baseline annotations.

**Supplementary Table 4: MS GWAS cell-specific enrichment within terminal hematopoietic cell types.** Columns are the same as in Supplementary Table 1.

**Supplementary Table 5: Heritability enrichments (single model) for other disorders.** Heritability enrichment p-values for each cell types across 10 autoimmune or neuropsychiatric disorders.

**Supplementary Table 6: Heritability enrichments (joint model) for other disorders.**

Heritability enrichments (p-values for coefficient *τ_c_*) for each cell type under joint model across 10 autoimmune or neuropsychiatric disorders.

**Supplementary Table 7. MS GWAS enrichment in CD4+ T cell subpopulations.** Columns are the same as in Supplementary Table 1.

**Supplementary Table 8. Stratified LDSC in CD4+ T cell subpopulations.** Columns are the same as in Supplementary Table 2.

**Supplementary Table 9.** MS GWAS pairwise enrichment in CD4+ T subpopulations.

Columns are the same as in Supplementary Table 3.

**Supplementary Table 10. MS GWAS enrichment in B cell subpopulations.** Columns are the same as in Supplementary Table 1.

**Supplementary Table 11. Stratified LDSC in B cell subpopulations.** Columns are the same as in Supplementary Table 2.

**Supplementary Table 12. MS GWAS pairwise enrichment in B cell subpopulations.**

Columns are the same as in Supplementary Table 3.

**Supplementary Table 13. Clinical characteristics of MS subjects.** EDSS: Expanded Disability Status Scale. * for the Untreated subjects it indicates number of months since last treatment.

**Supplementary Table 14. MS GWAS enrichment across OCRs in 6 cell types isolated from untreated MS patients.** Columns are the same as in Supplementary Table 1.

**Supplementary Table 15. MS GWAS enrichment across OCRs in joint model in CD4 T cell types isolated from untreated MS patients.** Columns are the same as in Supplementary Table 2.

**Supplementary Table 16. MS GWAS enrichment across OCRs in joint model in B cell types isolated from untreated MS patients.** Columns are the same as in Supplementary Table 2.

**Supplementary Table 17 MS GWAS enrichment across OCRs in 6 cell types isolated from treated MS patients.** Columns are the same as in Supplementary Table 1, except column added for the immune-modulating treatment

**Supplementary Table 18. MS GWAS enrichment across OCRs in joint model in T4em isolated from untreated and treated MS patients.** Columns are the same as in Supplementary Table 2

**Supplementary Table 19. MS GWAS enrichment across OCRs in joint model in cMBc isolated from untreated and treated MS patients.** Columns are the same as in Supplementary Table 2

**Supplementary Table 20. MS GWAS enrichments in histone ChIP-seq from T_h_17 cells.**

Columns are the same as in Supplementary Table 1.

**Supplementary Table 21. MS GWAS enrichments in histone ChIP-seq from B cells.**

Columns are the same as in Supplementary Table 1.

**Supplementary Table 20. MS GWAS enrichments in chromatin states from T_h_17 cells.**

Columns are the same as in Supplementary Table 1.

**Supplementary Table 21. MS GWAS enrichments in chromatin states from B cells.**

Columns are the same as in Supplementary Table 1.

**Supplementary Table 24: Summary of gene prioritizations.** For each GWAS locus, the lead SNP (“Effect SNP”), region annotation according to Patsopoulos et al., lead SNP chromosome, lead SNP position (in hg19), A1 allele, A2 allele, odd’s ratio (OR) and lead SNP p-value are shown. Previous gene prioritizations from Patsopolous et al., based on various criteria are listed. For columns “B Cell ATAC (Buenrostro)”, “B Cell ATAC (Calderon)”, ”Memory B Cell ATAC (Calderon)”, ”CD4 Cell ATAC (Buenrostro)”, ”CD4 Cell ATAC (Calderon)”, ”Th17 Cell ATAC (Calderon)”, a “1” indicates that a credible set SNP in the locus directly intersects an OCR from the indicated ATAC-seq dataset (“0” if no credible set SNP in the locus directly intersects. “PCHiC CD4 genes” and “PCHiC B genes” columns list genes that have a promoter capture HiC looping interaction to a credible set SNP in the indicated cell type. “PCHiC + ATAC CD4 genes” indicates a credible set SNP intersect an OCR in CD4 T cells and forms a PCHiC looping interaction in CD4 T cells to that gene. “PCHiC + ATAC B genes” indicates a credible set SNP intersect an OCR in B cells and forms a PCHiC looping interaction in B cells to that gene. The number of credible set SNPs in each locus is shown in the last column.

**Supplementary Table 25: Canonical pathway enrichment for prioritized MS genes.**

**Supplementary Table 26: Protein-protein interaction connectivity summaries for prioritized gene lists.** Detailed outputs from GeNets are provide.

**Supplementary Table 27: CD4 T prioritized genes ranked in the opposite extreme 10% of gene expression changes in KD and OE cell lines.**

**Supplementary Table 28: B prioritized genes ranked in the opposite extreme 10% of gene expression changes in KD and OE cell lines.**

**Supplementary Table 29: Markers for FACS sorting strategy of Verily ATAC-seq data**

